# Random Tanglegram Partitions (Random TaPas): An Alexandrian Approach to the Cophylogenetic Gordian Knot

**DOI:** 10.1101/481846

**Authors:** Juan Antonio Balbuena, Óscar Alejandro Pérez-Escobar, Cristina Llopis-Belenguer, Isabel Blasco-Costa

**Author notes:** **Corresponding author:**, Juan A. Balbuena, Ecology and Evolution of Symbionts Lab, Cavanilles Institute of Biodiversity and Evolutionary Biology, University of Valencia, Official P.O. Box 22085, 46071, Valencia, Spain., Cell.

## Abstract

Symbiosis is a key driver of evolutionary novelty and ecological diversity, but our understanding of how macroevolutionary processes originate extant symbiotic associations is still very incomplete. Cophylogenetic tools are used to assess the congruence between the phylogenies of two groups of organisms related by extant associations. If phylogenetic congruence is higher than expected by chance, we conclude that there is cophylogenetic signal in the system under study. However, how to quantify cophylogenetic signal is still an open issue. We present a novel approach, Random Tanglegram Partitions (Random TaPas) that applies a given global-fit method to random partial tanglegrams of a fixed size to identify the associations, terminals and nodes that maximize phylogenetic congruence. By means of simulations, we show that the output value produced is inversely proportional to the number and proportion of cospeciation events employed to build simulated tanglegrams. In addition, with time-calibrated trees, Random TaPas is also efficient at distinguishing cospeciation from pseudocospeciation. Random TaPas can handle large tanglegrams in affordable computational time and incorporates phylogenetic uncertainty in the analyses. We demonstrate its application with two real examples: Passerine birds and their feather mites, and orchids and bee pollinators. In both systems, Random TaPas revealed low cophylogenetic signal, but mapping its variation onto the tanglegram pointed to two different coevolutionary processes. We suggest that the recursive partitioning of the tanglegram buffers the effect of phylogenetic nonindependence occurring in current global-fit methods and therefore Random TaPas is more reliable than regular global-fit methods to identify host-symbiont associations that contribute most to cophylogenetic signal. Random TaPas can be implemented in the public-domain statistical software R with scripts provided herein. A User’s Guide is also available at GitHub.

---

Symbiosis is widespread throughout the tree of life and is considered as a key driver of evolutionary novelty and ecological diversity (Moran 2006; Zook 2015). Because organisms do not evolve in isolation, the evolutionary fate of symbiotic partners is intertwined at ecological and evolutionary levels, but despite the centrality of symbiosis in evolutionary biology, our understanding of how macroevolutionary processes originate extant symbiotic associations is still very incomplete (Weber et al. 2017). However, the recent emergence of robust comparative phylogenetic methods has expanded and facilitated research in this area (Hutchinson et al. 2017a). Cophylogeny, in particular, provides a quantitative framework to evaluate the dependency of two evolutionary histories (Hutchinson et al. 2017b). This approach involves some assessment of the congruence between the phylogenies of two groups of species or taxa related by extant associations, where congruence quantifies the degree of both topological and branch-length similarity (Page 2003). If such congruence is higher than expected by chance, it is concluded that there is cophylogenetic signal in the system studied (Mendlová et al. 2012).

Although cophylogenetic signal was initially interpreted as evidence of high level of cospeciation, it has been shown that other mechanisms can account for some degree of topological congruence (Kahnt et al. 2019). Particularly, complete host-switching events (i.e., colonization of a new host species followed by speciation) among closely related hosts can result in symbiont diversification mimicking the tree topology of the host, a process that has been termed preferential host-switching (Charleston and Robertson 2002) or pseudocospeciation (de Vienne et al. 2013). In any case, even if its causality cannot not be determined, quantifying cophylogenetic signal is highly relevant because it implies that contemporary ecological associations among species have been the product of a coupled evolutionary history.

The wide range of cophylogenetic methods currently available can be roughly categorized as either event-based or global-fit (Hutchinson et al. 2017a), but in our opinion none of them quantifies cophylogenetic signal satisfactorily. Event-based methods attempt to reconstruct the coevolutionary history of the organisms involved by assigning costs to each type of event and heuristically search for the solution(s) that minimize(s) the overall sum of costs (Charleston and Libeskind-Hadas 2014). The problem is that cophylogenetic signal can be overestimated because the default cost of cospeciation is assumed to be strictly less than the other events, which is often at odds with empirical evidence (de Vienne et al. 2013). In addition, with large datasets, the approach becomes computationally prohibitive and the influence of phylogenetic uncertainty is not explicitly considered (i.e., the single input phylogenetic trees of host and symbionts are assumed to represent the actual evolutionary relationships), which may lead to erroneous conclusions if not all clades are well supported. Global-fit methods, for their part, assess the degree of congruence between two phylogenies and can also identify the specific interactions that contribute most to overall congruence (Balbuena et al. 2013). They can handle large datasets economically in terms of computational time, and the effect of phylogenetic uncertainty can be assessed (Pérez-Escobar et al. 2016). However, current methods such as PACo (Balbuena et al. 2013) or ParaFit (Legendre et al. 2002), provide statistical evidence of cophylogenetic signal, but produce no clearly interpretable statistic as for its strength and no explicit links with coevolutionary events are made.

Therefore, additional work remains to be done in this domain. Particularly, the recent, spectacular expansion of DNA sequencing and phylogenetic reconstruction is leading to increasingly common scenarios involving species-rich trees and complex host-symbiont interactions (e.g., Hutchinson et al. 2017b). Envisaging these elaborate systems as evolutionary Gordian knots, we present herein an Alexandrian approach to cophylogenetic signal assessment. Random Tanglegram Partitions (Random TaPas) applies a given global-fit method to random partial tanglegrams of a fixed size to identify the associations, terminals and nodes that maximize phylogenetic congruence. By means of simulations, we show that the output value produced by Random TaPas is inversely proportional to both the number and proportion of cospeciation events to the total number of coevolutionary events employed to build the tanglegrams. In addition, with time-calibrated trees, Random TaPas is also efficient at distinguishing cospeciation from pseudocospeciation. The method is also useful to identify what host-symbiont associations contribute most to phylogenetic congruence because the variation in cophylogenetic signal can be mapped onto the tanglegram and phylogenetic uncertainty can be incorporated in the analyses. We illustrate its application with two real datasets of Passerine birds and their feather mites (Klimov et al. 2017) and Neotropical orchids and their euglossine bee pollinators. R scripts (R Core Team 2018) to perform Random TaPas and a User’s Guide based on data from Lagrue et al. (2016) is also available at GitHub (https://github.com/Ligophorus/RandomTaPas).

## Materials and methods

### The Random-TaPas algorithm

The starting point is a triple (*H, S*, **A**), where *H* and *S* represent the phylogenies of hosts and symbionts, and **A** is a binary matrix with rows and columns corresponding to terminals in *H* and S, respectively, in which extant associations between each terminal are coded as 1 and no associations as 0. The triple is often represented graphically as a tanglegram (Fig. 1), in which *H* and *S* usually are displayed face to face and their terminals are connected by lines reflecting the associations encapsulated in **A**. Figure 1 provides a complete overview of the Random TaPas algorithm. In short, for a large number of times (*N*), it selects *n* random unique one-to-one associations, so that each taxon in *H* is associated to one, and only one, taxon in *S*, and vice versa. (Since the aim is to find associations that maximize congruence, this is a condition to be met by any perfect host-symbiont cospeciation scenario). Then a global-fit method is applied to the partial tanglegram defined by the *n* associations and the statistic produced is saved. Finally, a percentile *p* of the distribution of the *N* statistics generated is set and the frequency of occurrence of each host-symbiont association of **A** included in *p* is computed.

**Figure 1.**
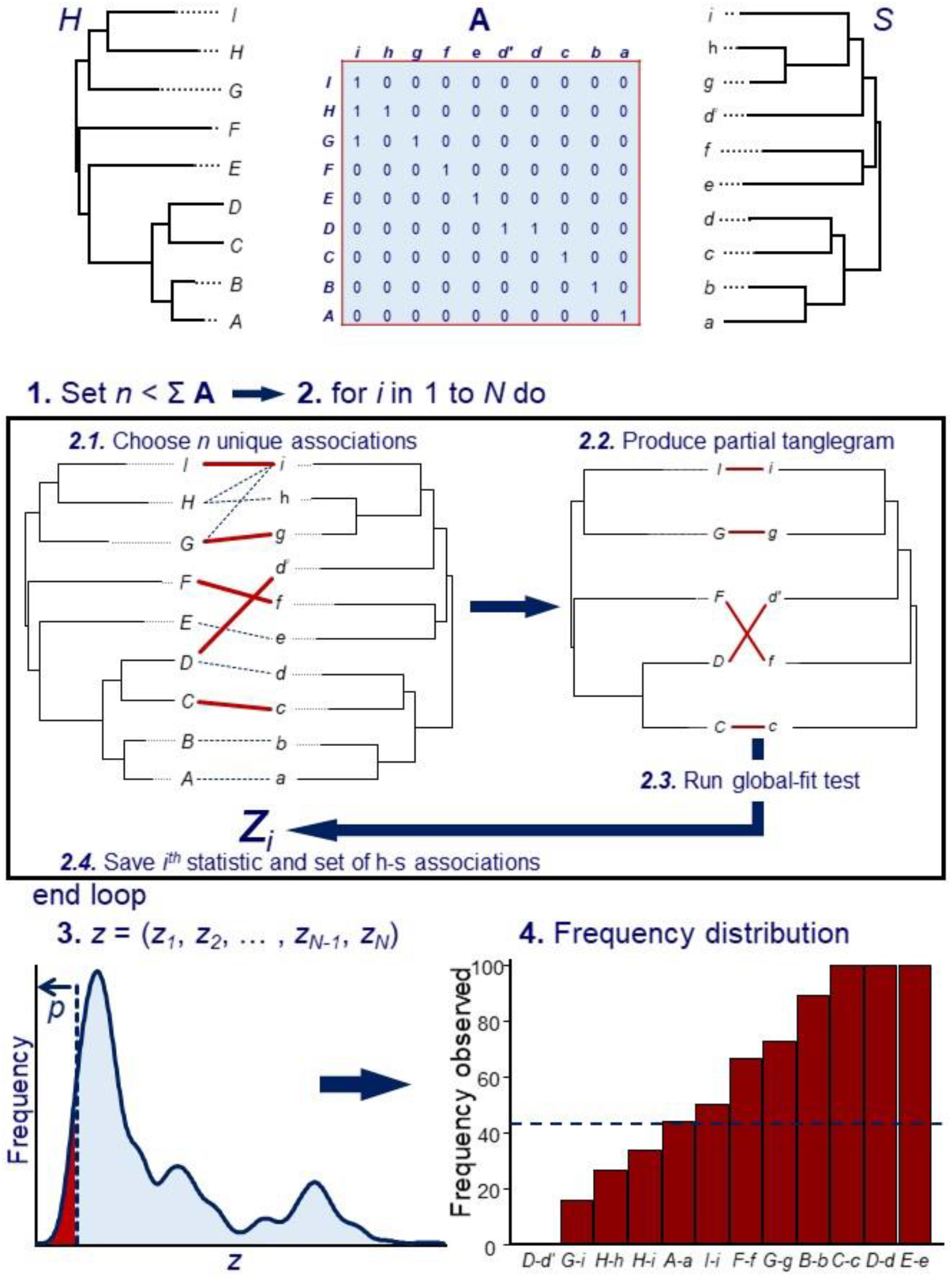
The Random TaPas algorithm. Given a triple (*H, S*, **A**), where *H* and *S* represent the phylogenies of hosts and symbionts, and **A** is a binary matrix that codes the associations between terminals in *H* and *S*: **1.** Set a number *n* less than the total number of host-symbiont associations. **2.** For *i* from 1 to *N* times (where *N* is sufficiently large, typically ≥ 10^4^) do ***2.1.*** Randomly choose *n* unique associations in **A**, so that each terminal in *H* is associated to one, and only one, terminal in *S*, and vice versa. (This is a prerequisite for a perfect host-symbiont cospeciation scenario); ***2.2.*** Produce a partial tanglegram that includes only the *n* associations chosen at step 2.1 by trimming A and pruning *H* and *S*; ***2.3.*** Run a global-fit test with the partial triple and ***2.4.*** Save the resulting statistic *z_i_* and the set of *n* host-symbiont associations selected at 2.1. **3.** Render the frequency distribution of the *z_i_*’s and set a small percentile *p* where the highest cophylogenetic congruence is expected. (In this example, *z_i_* is expected to be inversely proportional to congruence). **4.** Determine how many times each host-symbiont associations occurs in *p*, and return their frequency distribution.

The output of Random TaPas is a frequency distribution of host-symbiont associations (Fig. 1). Initially each value is expected to reflect the contribution of individual host-parasite associations to the global pattern of phylogenetic congruence. However, since the sampling scheme of Random TaPas favors the selection of one-to-one associations over multiple ones, it can lead to the former being overrepresented in the overall frequency distribution and hence in the percentile *p*. So their relative contribution to congruence could be overestimated, especially if the number of multiple associations is high. This bias can be corrected by determining the frequency of each host-symbiont association in the whole frequency distribution over the *N* runs (i.e., the distribution depicted at Step 3, Fig. 1). Assuming a null model in which the occurrence of each host-symbiont association is evenly distributed along the whole frequency distribution, the expected frequency of link *i* in percentile *p* (*Ep_i_*), can be estimated as *Ep_i_* = *n_i_* · *p*, where *n_i_* is the number of occurrences of the *i*^th^ association in the whole frequency distribution, and *p* is expressed as a decimal. Under the null model, the observed frequency of link *i* in percentile *p* (*Op_i_*) should equal *Ep_i_*, so the residual *Rp_i_* = *Op_i_* – *Ep_i_* would indicate whether link *i* is represented more often than expected by chance in *p*. Thus, in tanglegrams with a large portion of multiple host-symbiont associations, the distribution of residuals should be preferred as output over the initial frequency distribution of host-symbiont associations.

The shape of distributions can be characterized by means of the Gini coefficient (*G*), which is a common measure of inequality of a distribution (Ultsch and Lötsch 2017). When all distribution values are positive, the Gini coefficient is bounded between 0 and 1, representing respectively minimal and maximal inequality of the distribution. However, the inclusion of negative values, as it occurs in the distribution of residuals produced by Random TaPas, can lead to coefficients outside that range. Therefore, we resorted to measure dispersion of the distribution of residuals using the normalized Gini coefficient (*G*\*) devised by Raffinetti et al. (2015) to keep the values within the 0-1 interval. The computation of *G*\* is identical to that if the conventional Gini coefficient except that the arithmetic mean of observations is replaced by a normalized term to take account of the negative observations. Thus *G*\* can be expressed as

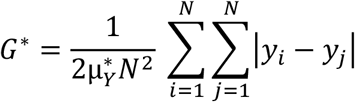

where *N* is the number of host-symbiont associations, *y_i_* and *y_j_* represent elements of the vector of residuals *Y* = (*y*_1_, *y*_2_, …, *y_N_*), and 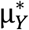 is the normalized term computed as 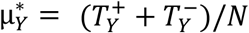, where 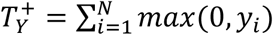 and 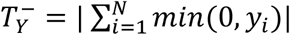 (i.e., the sum of positive residuals and absolute sum of negative residuals, respectively) (Raffinetti et al. 2015). Note that if all *y_i_* > 0, *G*\* = *G*.

We posit that the value of *G*\* in the present context is biologically informative because it is expected to be inversely proportional to cophylogenetic signal. In a perfect cospeciation scenario (maximal cophylogenetic signal), each host-symbiont association contributes equally to the global fit rendering *G*\* = 0. In contrast, extremely unequal contributions of the host-symbiont associations would yield highly skewed frequency distributions of residuals, in which *G*\* would approach one and cophylogenetic signal would be very small. It is also of interest to consider the scenario in which each host-symbiont association has equal chance to be associated with any residual value within the observed range. This can be modelled with a random uniform continuous distribution for which the expected value of the conventional Gini coefficient is 1/3 (Ultsch and Lötsch 2017). For *G*\*, if we assume that the expected distribution of residuals under the null model is centered at zero it can be proved that the expected value is ⅔ (See proof in Supplementary Material). So, ⅔ (or ⅓ in one-to-one-host-symbiont associations) can be initially taken as a threshold to determine whether a given tanglegram exhibits higher or lower cophylogenetic signal than expected by chance.

### Global-fit methods

Random TaPas can be applied in conjunction with any global-fit method. For the sake of demonstration, we chose here two very different approaches: Procrustes Approach to Cophylogeny (PACo) (Balbuena et al. 2013) and geodesic distances (GD) in tree space (Schardl et al. 2008). PACo uses Procrustean superimposition of Euclidean embeddings of the phylogenetic trees to assess phylogenetic congruence. The second method entails computing pairwise GDs between the partial phylogenies defined by the *n* associations. Most global-fit methods translate matrices of evolutionary distances into Euclidean space whose dimensionality is higher than that of tree space (Holmes 2005). So the advantage of this approach is that it skips the potential effect of this dimensional mismatch. In both methods, the value of the statistic produced is inversely proportional to topological congruence between the trees evaluated.

PACo was performed with package paco (Hutchinson et al. 2017b) in R, employing patristic distances as input. Its use with Random TaPas is aimed at finding the set of partial tanglegrams that minimize the square sum of residuals 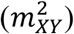 of the Procrustes superposition of host and symbiont spatial configurations in *p*. GDs were computed with function `dist.multiPhylo` of `package` distory in R (Chakerian and Holmes 2010) in order to find the set of partial tanglegrams in *p* with minimal pairwise distances in tree space between the host and symbiont partial phylogenies.

### Synthetic data

The operation of Random TaPas depends on three parameters: *N, n* and *p* (Fig. 1). We assessed performance with different parameter combinations and the two aforementioned global fit methods with 200 synthetic tanglegrams generated with CoRe-Gen (Keller-Schmidt et al. 2011). For a given set of input parameters, the program provides a pair of resulting ultrametric trees, a list of associations between the terminals and the number of coevolutionary events (cospeciation, duplication, sorting, and complete host-switching) involved in the construction of the tanglegram. Since it is not possible to accurately control a priori the output based on the input parameters, we first generated a library of 1,000 tanglegrams. The trees were built following a pure-speciation (Yule) model, which has been shown to describe adequately empirical phylogenetic trees (Hagen et al., 2015). For each host-symbiont pair, we specified a different random combination of input parameters sampled from a uniform distribution. The parameters and sampling ranges were: number of generations (100-200), probability of cospeciation event (0.2-1.0), probability of host-switching event (0.2-0.8) and probability of choosing a host for speciation instead of a parasite (0.7-1.0). Because our method is intended for a wider range of settings than those considered by Keller-Schmidt et al. (2011), the parameter ranges are broader than those used originally. Thus, our resulting tanglegrams included scenarios where cospeciation events ranged from very rare to very common. Since some combinations of parameters yielded trees with few terminals (≤ 15) or numbers of host parasite associations (≤ 20), we ran CoRe-Gen 1,274 times to obtain a set of 1,000 workable tanglegrams.

Given that Random TaPas is specially intended for large tanglegrams, we chose for subsequent analyses two sets of 100 tanglegrams each involving approximately 50 (mean = 50.5, median = 51, range: 47-54) and 100 (mean = 99.7, median = 100, range: 93-107) host-symbiont associations, respectively. For convenience, these subsets will be henceforth referred to as Set50 and Set100, respectively. Since some trees included terminal polytomies and GD requires fully bifurcating trees, we added 0.01 arbitrary units of time to all terminal branches. A limitation of CoRe-Gen is that it only generates coevolutionary systems in which each symbiont can be associated with only a single host. Given that many real-world scenarios do not conform to this scenario, we modeled the occurrence of colonization (aka host-sharing) by a given symbiont of several hosts following Drinkwater et al. (2016). For each triple, a rate of host sharing of 0%, 10% or 20% of the total available host species was set randomly. Then colonization was simulated by selecting each terminal in *S* and allowing the corresponding symbiont to colonize a random number x of additional hosts, {*x* ∈ ℕ | 0 ≤ *x* ≤ *R* × *N_h_* }, where *R* represents the rate of host sharing for the given tanglegram and *N_h_* the total number of host terminals.

Since additive trees (i.e., phylograms) estimated from molecular sequence data are commonly used in cophylogenetic analyses, we ran the simulations with the original ultrametric trees and phylograms derived from them. The transformation of the ultrametric trees into phylograms was done by multiplying their branch lengths by varying rates of molecular substitution (Brown and Yang 2011; Paradis 2014), sampled from a log-normal distribution with mean 0.01 substitutions/site/time unit (Brown and Yang 2011).

### Relationship of G* with coevolutionary events

We applied Random TaPas to Set50 and Set100 (both additive and ultrametric trees) using both PACo and GD with all combinations of the following parameter values: *N*=10^4^, *n*= 5, 10 and 20 (Set50), and *n*=10, 20 and 40 (Set100) (i.e., *n* representing ≈ 10%, 20% and 40% of the total associations), and *p*= 1% and 5%. In preliminary analyses, *N*= 10^5^ was also tested with additive trees in combination with the same arrays of *n* and *p* values but the results were similar to those obtained with *N*=10^4^ (Supplementary Figs. S1 and S2 in Supplementary Material). Since the increased computing time resulting from using a larger *N* did not result in a detectable improvement in performance of the method, *N*= 10^5^ was not further considered. In simulations involving additive trees, prior to running PACo the patristic distances were rendered Euclidean by taking the element-wise square root of each cell in order to minimize geometric distortion and avoid negative eigenvalues (de Vienne et al. 2011). The frequency distributions obtained from each simulation were converted into distributions of residuals *Rp_i_* (observed – expected frequencies) as explained above.

For each simulation, the normalized Gini coefficient (*G*\*) of the distribution of residuals of host-symbiont associations produced by Random TaPas was computed with function `Gini_RSV` of package `GiniWegNeg` in R (Raffinetti and Aimar 2016). We assessed the relationship between *G*\* of each set of simulations, the number of coevolutionary events (cospeciation, duplication, sorting, host-switching and colonization) and the ratio of number of cospeciation events to the total number of coevolutionary events taken to build each tanglegram using Pearson’s correlation coefficients.

### Pseudocospeciation experiment

In order to assess the ability of Random TaPas to distinguish between cospeciation and pseudocospeciation (de Vienne et al. 2007), we selected a triple (#63) in Set50 with high cophylogenetic signal. This simulated host-symbiont system resulted from 50 cospeciations, 1 sorting, 2 duplications, 1 complete host-switching and 0 colonization events. We arbitrarily selected a clade of 17 terminals in the symbiont tree to simulate rapid colonization and speciation (de Vienne et al. 2007). In both the ultrametric and additive trees versions of the tanglegram, the branch lengths of the clade were shortened to half their original length, whereas its basal branch was lengthened to keep the original height of the clade. We applied Random TaPas to the original and modified tanglegrams using GD and PACo, *N* = 10^4^, *p* = 1%, and *n* = 5, 10 and 20. The ability of Random TaPas to distinguish cospeciation from pseudocospeciation was assessed by plotting as a heatmap on the tanglegram, the residual frequency of occurrence of each host-symbiont link and the average residual frequency of occurrence of each terminal in *p*.

### Mapping of congruent/incongruent associations

We also used simulated tanglegrams to evaluate the capacity of Random TaPas to map congruent and incongruent host-symbiont associations. We chose triple #84 in Set50 characterized by low cophylogenetic signal as this system resulted from 9 cospeciation, 20 sorting, 24 duplication, 10 complete host-switching and 4 colonization events. Then the same clade of 17 symbiont terminals used in the preceding cospeciation experiment and its corresponding clade of 16 host terminals of triple #63 in Set50 were inserted respectively in the host and symbiont trees of triple #84. The inserted clades represented an almost perfect cospeciation pattern except from one symbiont that for this simulation was associated randomly to a host (H5) of triple #84. We performed three different types of simulations in which clade insertions were implemented (1) at the base, (2) at central nodes (29 and 59 of the host and symbiont trees, respectively), and (3) at upper nodes (31 and 86, respectively) of the corresponding #84 ultrametric and additive trees. We applied Random TaPas using GD and PACo, *N* = 10^4^, *p* = 1%, and *n* = 7 and 14 (which represented about 5% and 10% of the total number of tanglegram associations). The residual frequency of occurrence was plotted as a heatmap on the tanglegram, of each host-symbiont association and the average residual frequency of occurrence of each terminal in p. In addition, taking the latter as a continuous trait, fast maximum likelihood estimators of ancestral states were computed with fastAnc of package phytools in R (Revell 2012), and their values were displayed on the nodes of the phylogeny based on the same heatmap scale. We adopted this approach only to assess different levels of cophylogenetic signal across the tanglegram and by no means imply that we consider these estimators to reflect actual ancestral states of host-symbiont associations.

### Case Study Examples

To gain further insight on its performance and provide further guidance to prospective users, we demonstrate the application of Random TaPas to two real-world examples.

#### Example 1: Passerine birds and feather mites

We examined cophylogenetic patterns between passerine birds and proctophyllodid feather mites based on a published dated phylogeny of proctophyllodid feather mites and 200 Bayesian chronograms of their passerine hosts (Klimov et al. 2017).

Random TaPas was run with GD and PACo, *N* = 10^4^, *n* = 20 and *p* = 1% to evaluate the agreement between the passerine and mite evolutionary histories. The *n* chosen it is close to the 20% of the number of associations (98), which based on results of the preceding simulations seems to represent a good compromise between assessing cophylogenetic signal and detecting congruent host-symbiont associations. For each host tree, a separate analysis was carried out with each global-fit method and then the average residual frequency per bird-mite association of the 200 runs was mapped on the tanglegram. (We randomly chose a host chronogram to plot the tanglegram). In addition, the average residual frequency of occurrence of each terminal occurring in p, and ancestral states were estimated and mapped on the tanglegram as per the preceding experiment.

#### Example 2: Neotropical orchids and their euglossine bee pollinators

In addition to the analysis of cophylogenetic relationships between hosts and symbionts, we demonstrate in this example how to assess the influence of phylogenetic uncertainty in the analyses. To this end, we used the association data and published chronograms of pollinated orchids and their corresponding bee pollinators (Ramírez et al. 2011), and the posterior probability trees used to build the respective consensus trees, which were kindly supplied by Santiago Ramírez, University of California Davis.

Random TaPas was run with GD and PACo, *N* = 10^4^, *n* = 26 (i.e., about 20% of the 129 bee-orchid associations) and *p*= 1%. In order to account for phylogenetic uncertainty, we computed 95% confidence intervals of the host-symbiont residual frequencies of each association using a random sample (excluding the burn-in set) of 1,000 pairs of the posterior-probability trees used to build the consensus trees of euglossine bees and orchids. Random TaPas was run for each tree pair as specified above and the 95% confidence interval of each host-symbiont association residual frequency were computed empirically based on the 1,000 runs.

We also assessed the different contribution of each host-symbiont association to the global cophylogenetic signal by displaying as a heatmap on the tanglegram their residual frequency of occurrence and the average residual frequency of occurrence of each terminal occurring in *p*, and estimators of ancestral states as described in Mapping of congruent/incongruent associations above.

## Results

### Relationship of G* with coevolutionary events

Not all simulations could be run because, in some triples, the number of possible unique one-to-one associations between terminals was less than the number *n* set in simulations. So the number of triples produced in Set50 with *n* = 20, was 79; and those in Set100 with *n*=20 and *n*=40 was 99 and 63, respectively.

Forty-eight types of simulations were produced to analyze the relationship between the normalized Gini coeficients (*G*\*) of host-symbiont residual frequency distributions and the number of evolutionary events of tanglegrams involving the additive and the original ultrametric trees with all combinations of *n* and *p* and global-fit method employed. The mean *G*\*s of each simulation tended to be smaller in simulations involving ultrametric trees (range 0.605-0.719) than those obtained with additive trees (range 0.693-0.732) (Table S1 in Supplementary Material). Most notably the ranges of *G*\*s in the latter were smaller than those obtained in simulations with ultrametric trees. This occurred because the lowest-range values were higher in simulations performed with additive trees, whereas the highest-range values were similar in both types of simulations, as shown in Figure 2 for simulations performed with GD. (The corresponding simulations with PACo yielded very similar results and are displayed in Fig. S3, Supplementary Material.) Triple size had a slight effect on the values of *G*\*, since the means were slightly larger and the ranges slightly smaller in simulations involving Set100 (Table S1).

**Figure 2.**
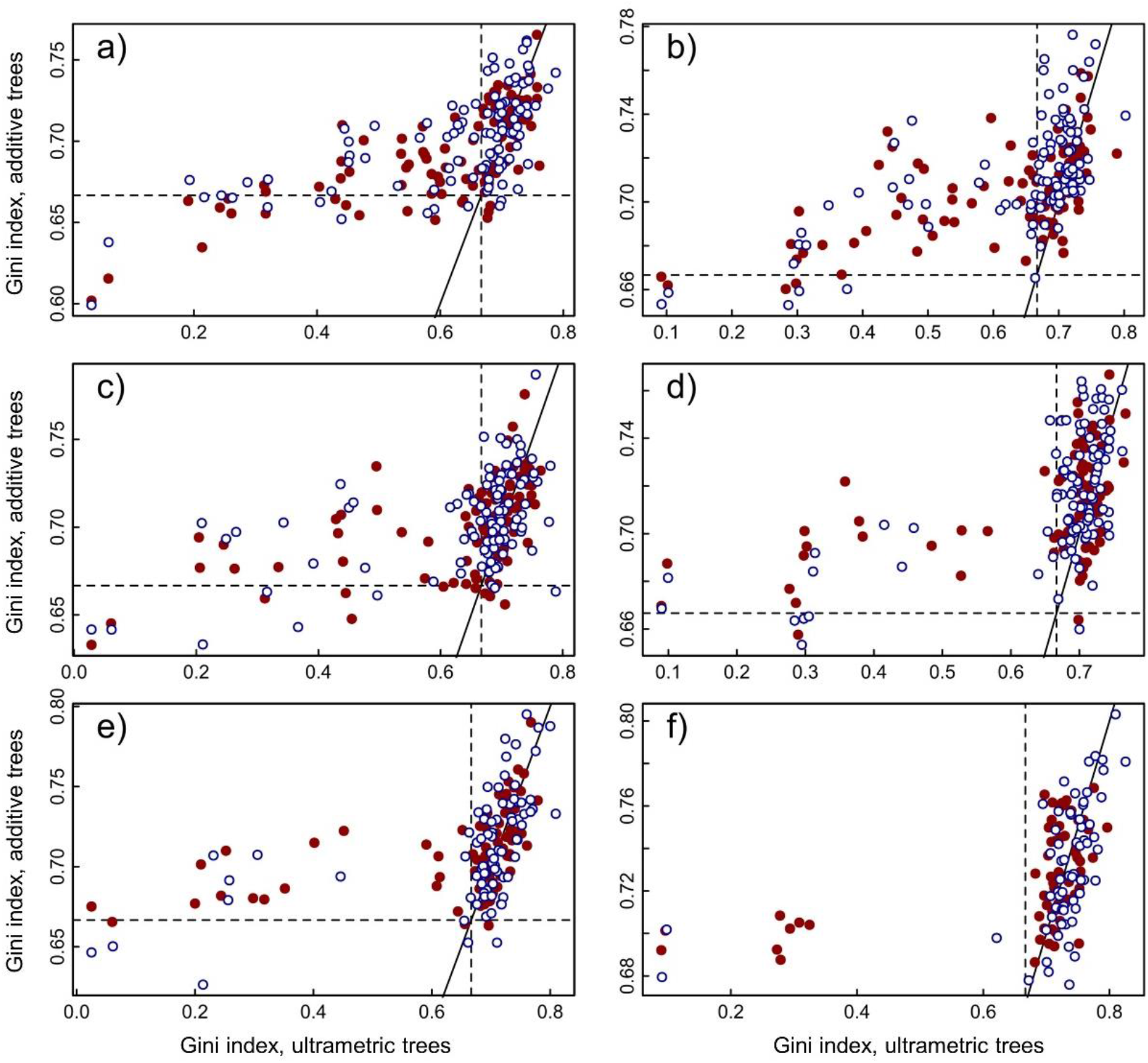
Comparison of the normalized Gini coefficient of the residual (observed – expected frequency) distribution of host-symbiont associations produced by Random TaPas using geodesic distances as global-fit method and ultrametric trees, with those based on additive trees. Parameter and test combinations: (a) Set50, *n* = 5; (b) Set100, *n* = 10; (c) Set50, *n* = 10; (d) Set100, *n* = 20; (e) Set50, *n* = 20; (f) Set100, *n* = 40. Filled red points, *p* = 1%; empty blue points, *p* = 5%. The dashed lines mark a theoretical threshold (⅔) between low and high cophylogenetic signal. The solid line represents *y* = *x*.

The correlation of *G\** with the number of cospeciation events was significant (*P* < 0.05) in all except two simulations involving Set100, additive trees, and GD as global-fit method (Fig. 3). The *G*\*s of the distribution frequency of residuals and the proportion of cospeciation events with respect to the total number of coevolutionary events was always significantly correlated and tended to be higher (in absolute value) than those involving the number of cospeciation events (Table 1). In both cases, the correlations with coevolutionary events and with the proportion of cospeciation events was stronger in tanglegrams based on ultrametric trees and in those of Set50. In addition, the highest absolute correlation values between *G*\* and the number and proportion of cospeciation events were observed with *p* = 1% and *n* representing about 5% of the total number of host-symbiont associations (Fig. 3, Table 1). This pattern was also observed when the number of three events causing incongruence, sorting, duplication and host-switching were considered. In contrast, the number of colonization events was usually weakly or not correlated with *G*\* (Fig. 3).

**Figure 3.**
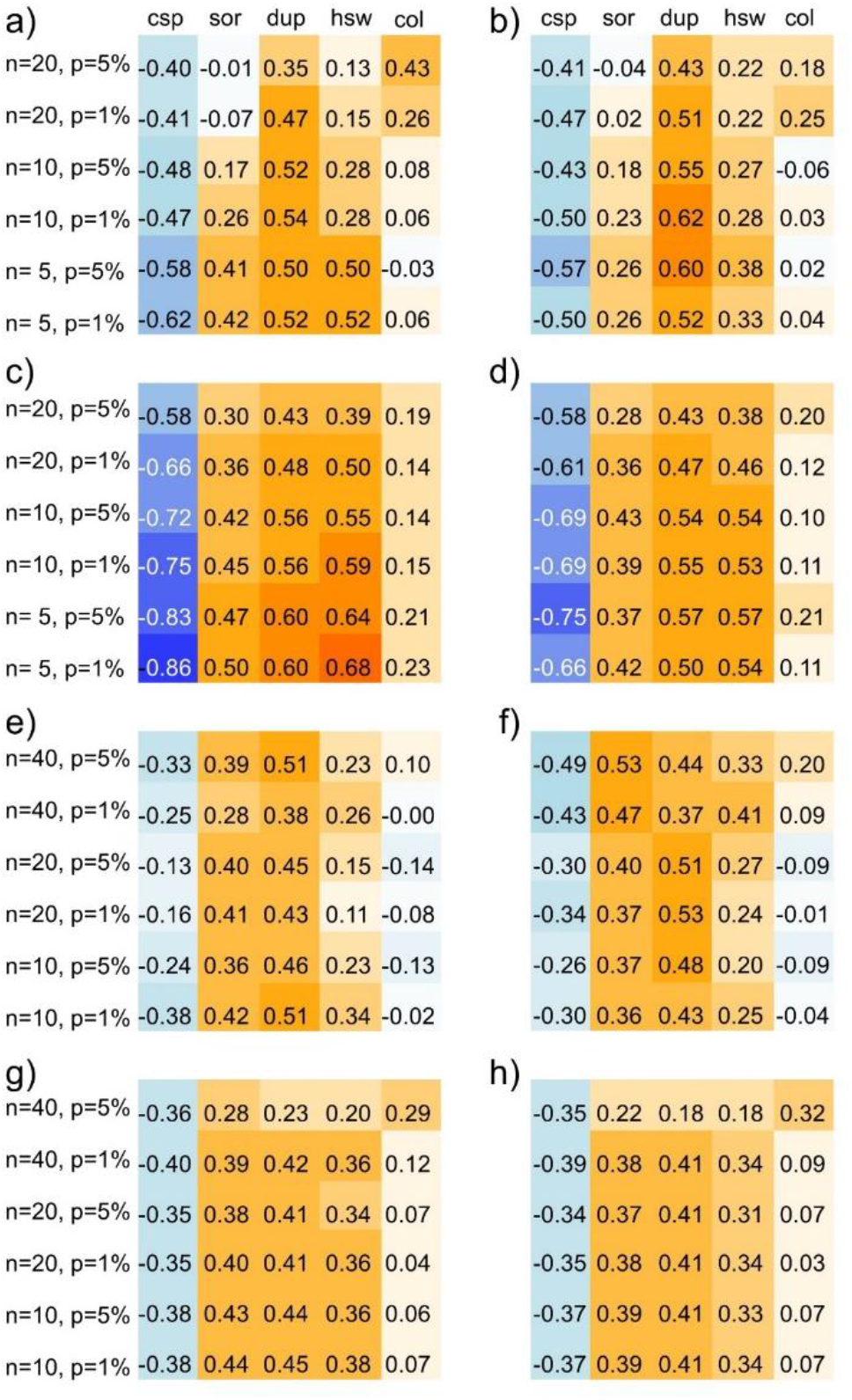
Correlation coefficients computed between the normalized Gini coefficients of the frequency distribution of residuals (observed – expected frequencies) of host-symbiont associations produced by Random TaPas and the number of cospeciation events in two sets of 100 simulated tanglegrams each involving approximately 50 (a-d) and 100 (e-h) host-symbiont associations, respectively. The tanglegrams were built with both additive (a, b, e, f) and ultrametric (c, d, g, h) trees and Random TaPas was applied with geodesic distances (a, c, e, g) and PACo (b, d, f, h) with a varying number of host-symbiont associations (n) and two percentile values (*p*). Event abbreviations: csp = cospeciation, sor = sorting, dup =duplication, hsw = complete host-switching, col = colonization of new host without speciation. Background colors indicate the strength of the correlation, ranging from blue (*r* =−1) to red (*r* =+1).

**Table 1:** Correlation coefficients of normalized Gini coefficients of the frequency distribution of residuals (observed – expected frequencies) of host-symbiont associations produced by Random TaPas with the proportion of cospeciation events with respect the total number of coevolutionary events in two sets of 100 simulated tanglegrams each involving ≈ 50 and 100 host-symbiont associations, respectively, and both additive (A) and ultrametric (U) trees applying Random TaPas with geodesic distances (GD) and PACo with a varying number of host-symbiont associations (*n*) and two percentile values (*p*). (See Fig. 1 for a definition of *n* and *p*).

| Tree | Set | $p$ | $n$ | GD | PACo |
| --- | --- | --- | --- | --- | --- |
| A | 50 | 1% | 5 | -0.65 | -0.50 |
| A | 50 | 5% | 5 | -0.60 | -0.57 |
| A | 50 | 1% | 10 | -0.49 | -0.50 |
| A | 50 | 5% | 10 | -0.47 | -0.43 |
| A | 50 | 1% | 20 | -0.38 | -0.47 |
| A | 50 | 5% | 20 | -0.38 | -0.40 |
| A | 100 | 1% | 10 | -0.44 | -0.35 |
| A | 100 | 5% | 10 | -0.31 | -0.33 |
| A | 100 | 1% | 20 | -0.26 | -0.40 |
| A | 100 | 5% | 20 | -0.26 | -0.38 |
| A | 100 | 1% | 40 | -0.34 | -0.51 |
| A | 100 | 5% | 40 | -0.44 | -0.57 |
| U | 50 | 1% | 5 | -0.89 | -0.70 |
| U | 50 | 5% | 5 | -0.86 | -0.78 |
| U | 50 | 1% | 10 | -0.79 | -0.73 |
| U | 50 | 5% | 10 | -0.76 | -0.73 |
| U | 50 | 1% | 20 | -0.69 | -0.65 |
| U | 50 | 5% | 20 | -0.60 | -0.59 |
| U | 100 | 1% | 10 | -0.50 | -0.45 |
| U | 100 | 5% | 10 | -0.50 | -0.47 |
| U | 100 | 1% | 20 | -0.48 | -0.45 |
| U | 100 | 5% | 20 | -0.47 | -0.45 |
| U | 100 | 1% | 40 | -0.52 | -0.49 |
| U | 100 | 5% | 40 | -0.43 | -0.40 |

### Pseudocospeciation experiment

Figures 4 and 5 show the results of the cospeciation experiments performed with GD. In the additive trees, host-symbiont associations involving the modified clade were marked with a similar level of congruence to those of the original tanglegram, independently of the *n* value chosen. In contrast, the counterpart associations in the ultrametric trees were marked as incongruent with *n* = 5 and 10, and partly (basal branches of the modified clade) with *n* =20. A similar pattern was observed in the corresponding experiments carried out with PACo (Figs. S4, S5 in Supplementary Material):

**Figure 4.**
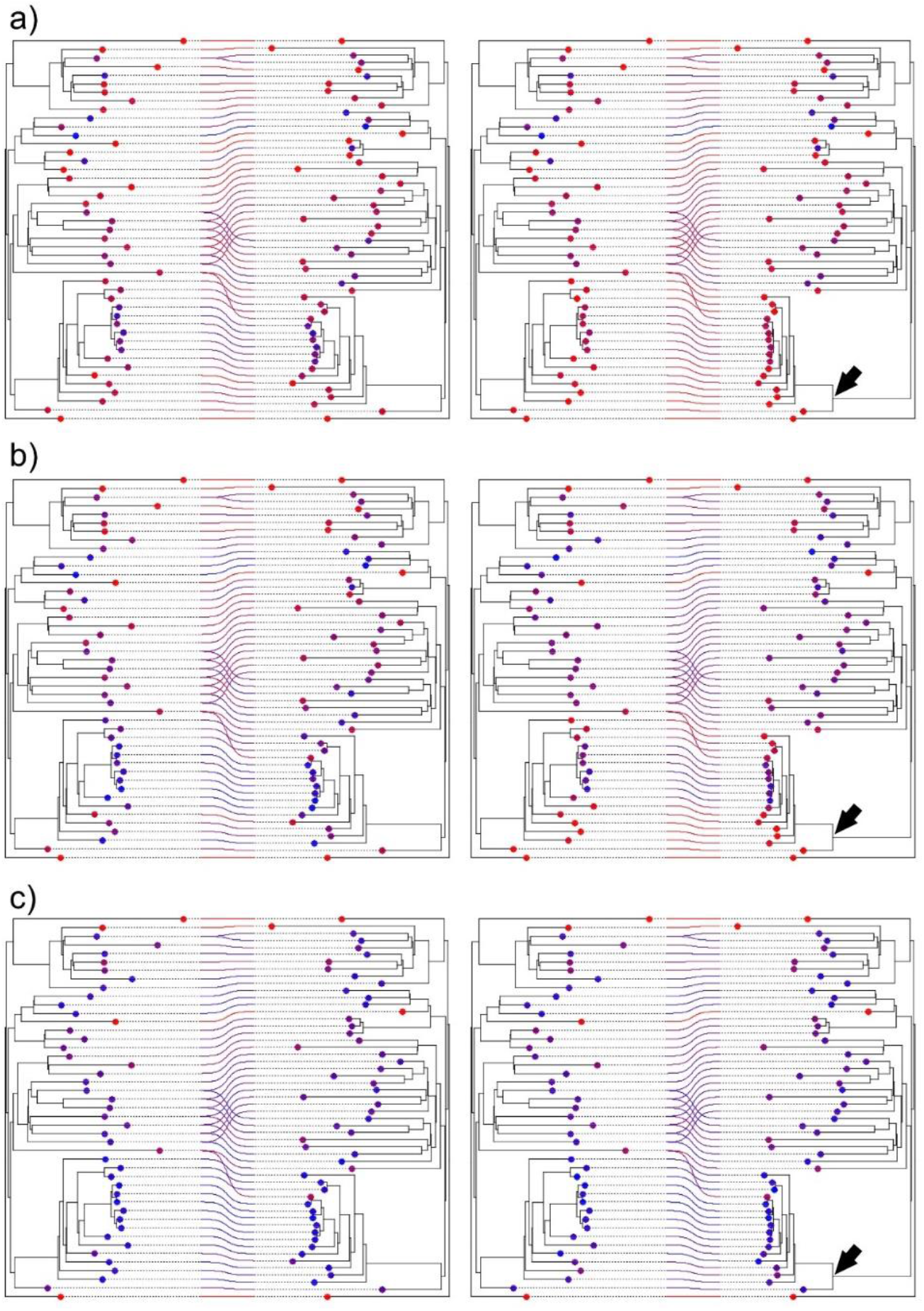
Pseudocospeciation experiment with one simulated tanglegram of ≈50 host-symbiont associations relating two additive trees. Random TaPas was applied with geodesic distances, *p* = 1% and *n* = 5 (a), *n* = 10 (b) and *n* = 20 (c) to the original (left) and modified (right) tanglegrams. In the latter, the branch lengths of one clade (arrow) were reduced to one half, whereas its basal branch was lengthened to keep the original height of the clade. The residual (difference between observed and expected frequency of occurrence of each host-symbiont association) in the percentile *p* retrieved by Random TaPas (see Fig. 1) is coded in a color scale, where red and blue denote low and high values, respectively. The points at terminals are also coded on the same scale and represent the average residual in which the terminal is involved.

**Figure 5.**
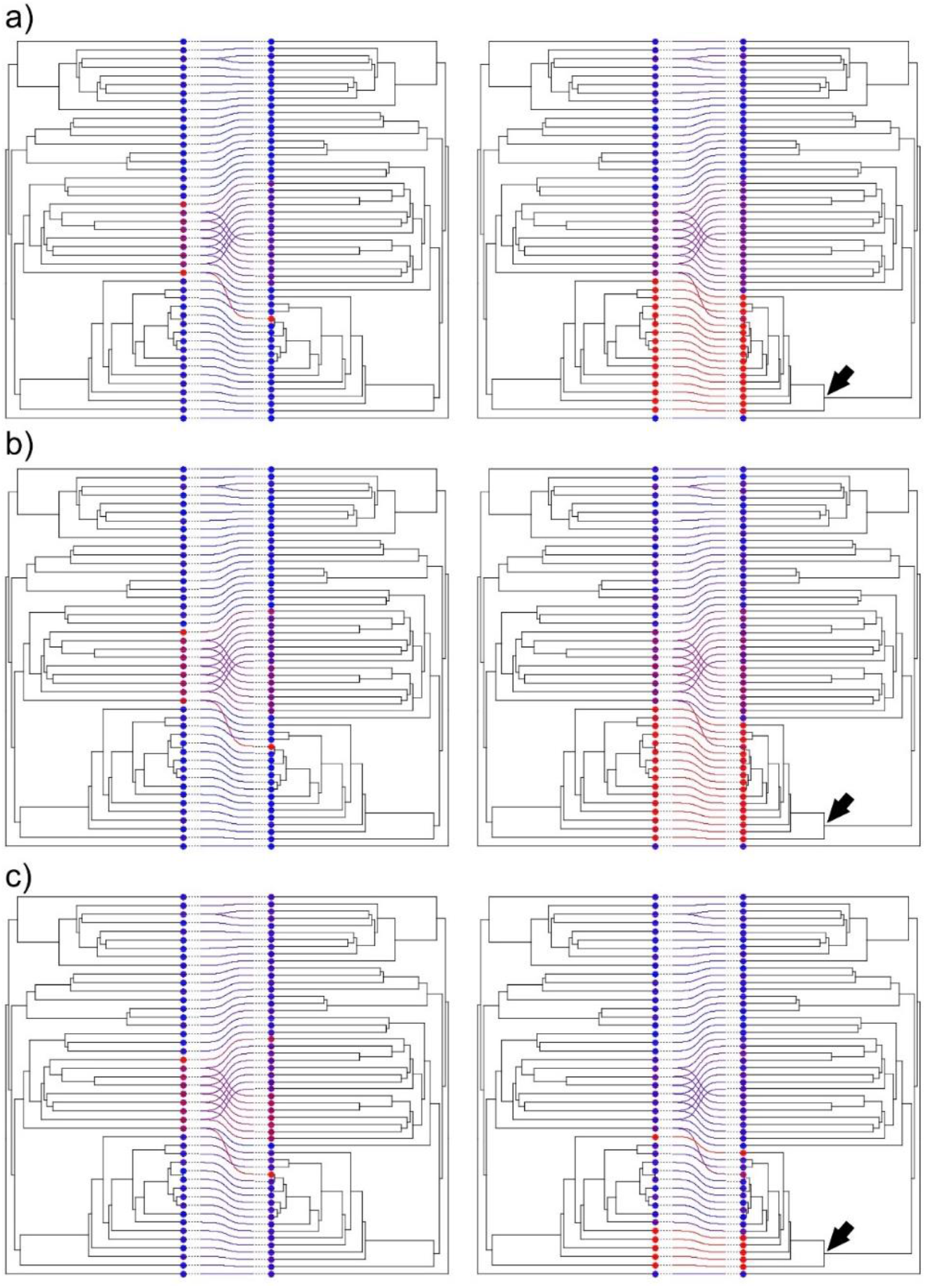
Pseudocospeciation experiment with one simulated tanglegram of ≈50 host-symbiont associations relating two ultrametric trees. Random TaPas was applied with geodesic distances, *p* = 1% and *n* = 5 (a), *n* = 10 (b) and *n* = 20 (c) to the original (left) and modified (right) tanglegrams. In the latter, the branch lengths of one clade (arrow) were reduced to one half, whereas its basal branch was lengthened to keep the original height of the clade. The residual (difference between observed and expected frequency of occurrence of each host-symbiont association) in the percentile *p* retrieved by Random TaPas (see Fig. 1) is coded in a color scale, where red and blue denote low and high values, respectively. The points at terminals are also coded on the same scale and represent the average residual in which the terminal is involved.

### Mapping of congruent/incongruent associations

Random TaPas was efficient at tagging the associations between the terminals of the inserted clades as relatively more congruent than those of the receptor tanglegram. This was particularly so with ultrametric trees and *n* = 14 (Figs. 6, 7 for GD and Figs. S6, S7 in Supplementary Material with PACo). With additive trees, some associations within the inserted clades were marked as incongruent, the degree of which seemed to be mostly determined by the absolute differences between branch lengths of the host and symbiont terminals involved (Fig. 6, Fig. S6).

**Figure 6.**
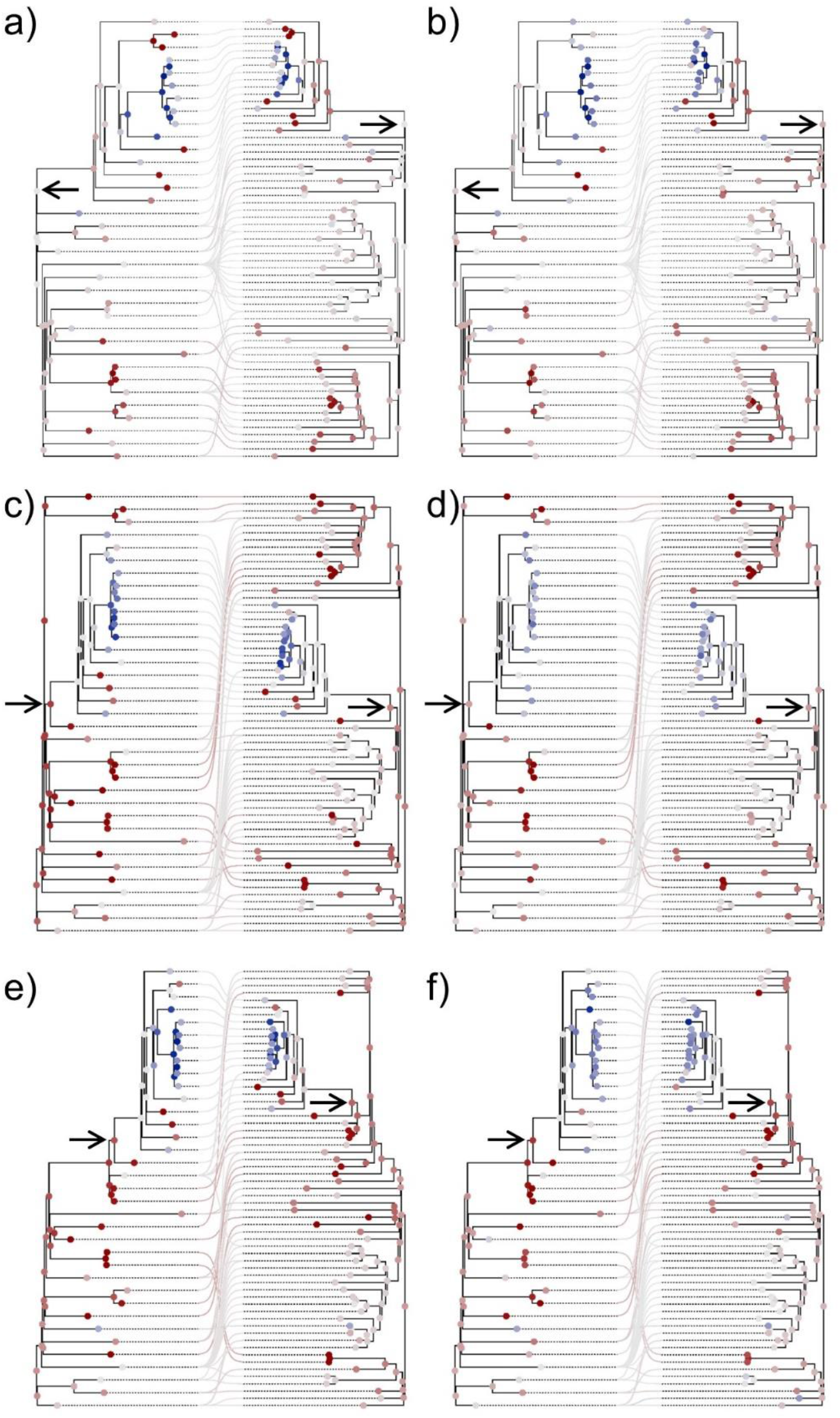
Simulated tanglegrams with additive trees in which a clade of host and symbionts (arrows) were inserted at different levels of the receptor tanglegram: at the root (a, b), and at middle (c, d) and upper (e, f) nodes. Random TaPas was applied with geodesic distances, *p* = 1% and *n* = 7 (a, c, e), *n* = 14 (b, d, f) to each tanglegram. The residual (difference between the observed and expected frequency of occurrence of each host-symbiont association) in the percentile *p* retrieved by Random TaPas (see Fig. 1) is mapped using a diverging color scale centered at zero (light gray) and ranging from dark red (maximum negative) to dark blue (maximum positive). The average observed-expected frequency of each terminal and fast maximum likelihood estimators of ancestral states of each node are also mapped according to the same scale.

**Figure 7.**
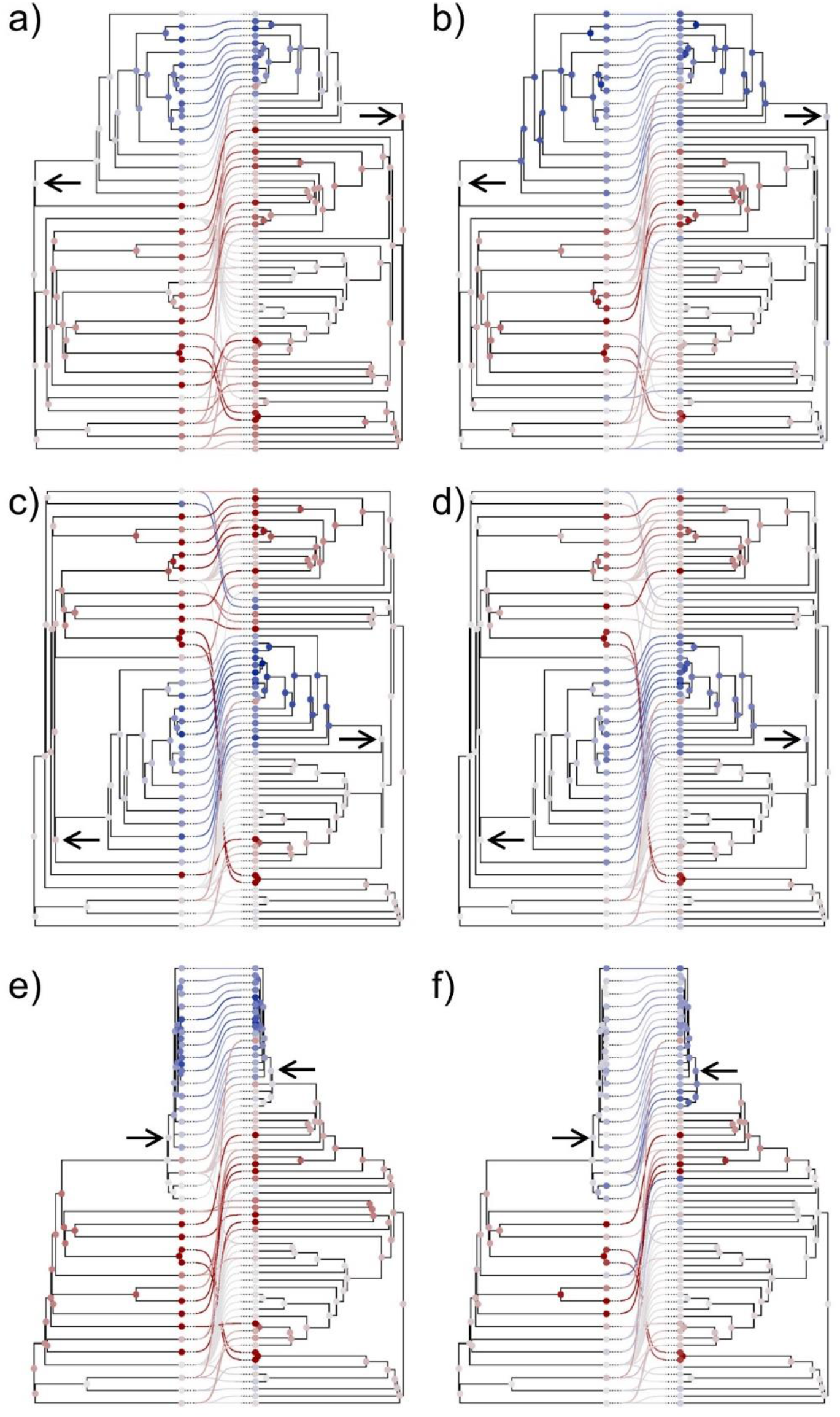
Simulated tanglegrams with ultrametric trees in which a clade of host and symbionts (arrows) were inserted at different levels of the receptor: at the root (a, b), and at middle (c, d) and upper (e,f) nodes. Random TaPas was applied with geodesic distances, *p* = 1% and *n* = 7 (a, c, e), *n* = 14 (b, d, f) to each tanglegram. The residual (difference between the observed and expected frequency of occurrence of each host-symbiont association) in the percentile *p* retrieved by Random TaPas (see Fig. 1) is mapped using a diverging color scale centered at zero (light gray) and ranging from dark red (maximum negative) to dark blue (maximum positive). The average observed-expected frequency of each terminal and fast maximum likelihood estimators of ancestral states of each node are also mapped according to the same scale.

### Example 1: Passerine birds and feather mites

Random TaPas applied to the 200 bird and 1 mite chronogram yielded 200 residual (observed – expected frequency) distributions for each of the comparisons performed with GD and PACo. Figure 9 displays boxplots of the *G*\*s associated to each distribution. The *G*\* values produced with GD and PACo were larger than the ⅔ threshold proposed to indicate random contributions of congruence by the host-symbiont associations.

**Figure 8.**
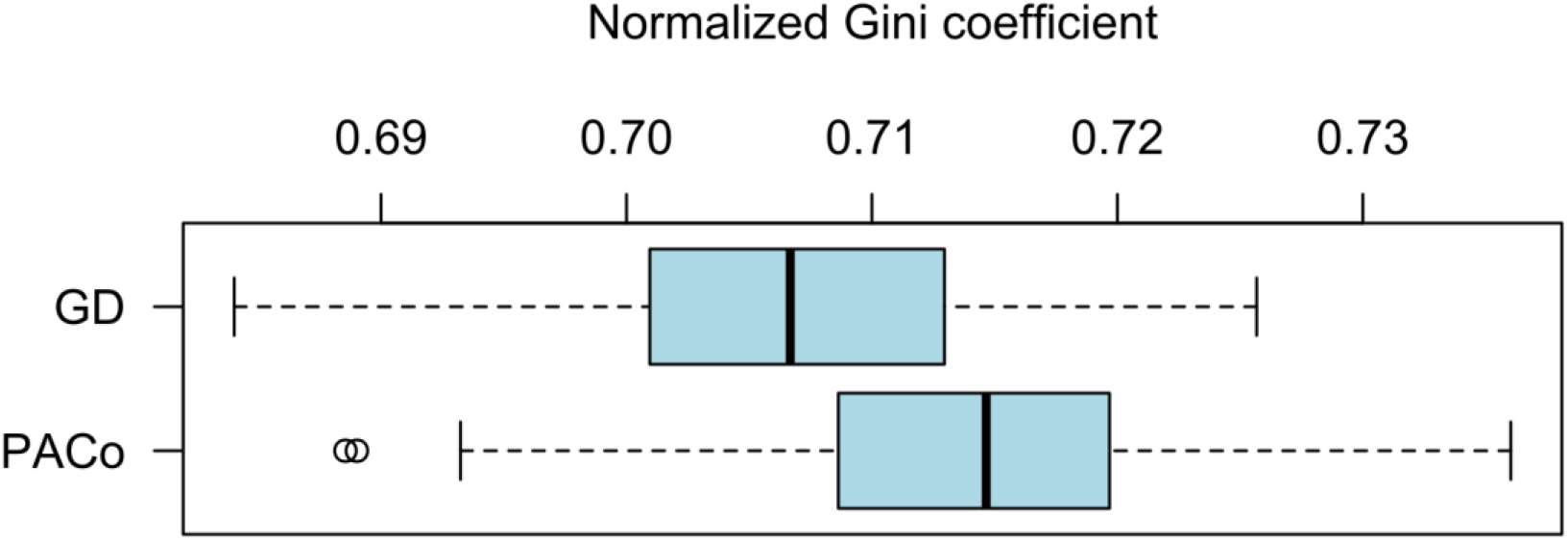
Boxplot summarizing the variation of 200 Normalized Gini coefficients of the respective residual frequency distributions produced with Random TaPas applied to pairs formed by 200 passerine bird chronograms and one proctophylloid mite chronogram using both GD and PACo as global fit methods.

**Figure 9.**
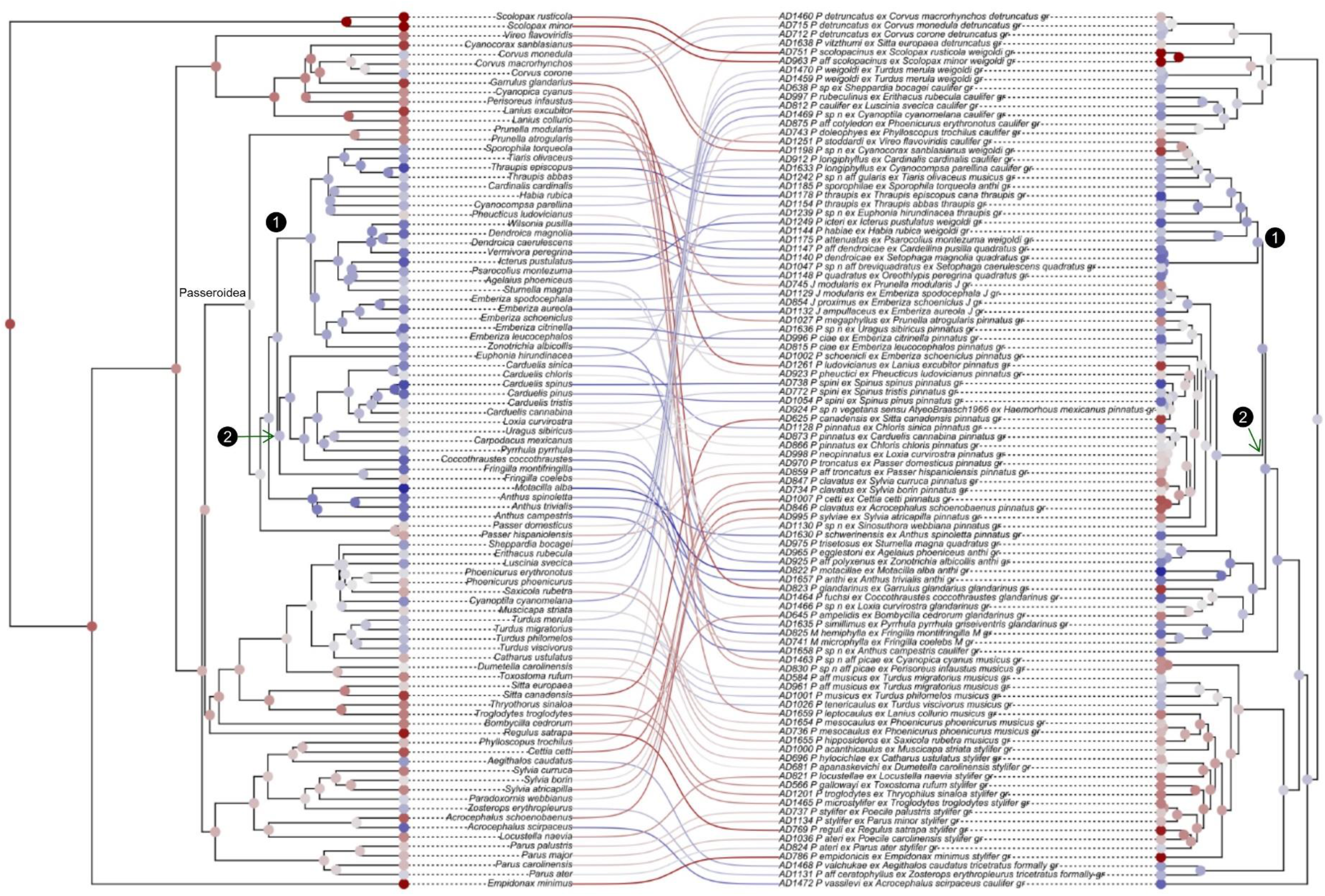
Tanglegram representing the association between passerine birds and their associated proctophyllodid feather mites. Random TaPas with GD was applied to 200 bird chronograms and one mite dated tree. For each host tree, a separate analysis was performed yielding a vector of residuals (observed-expected frequencies). The average residual over the 200 runs corresponding to each bird-mite association is mapped using a diverging color scale centered at zero (light gray) and ranging from dark red (maximum incongruence) to dark blue (maximum congruence). The average residual at each terminal and fast maximum likelihood estimators of ancestral states of each node are also mapped according to the same scale. Based on Klimov et al. (2017), two synchronic events are also indicated on the host and mite chronograms: (1) Diversifications of New World emberizoid Passerida and the *Proctophyllodes thraupis+quadratus* clade and (2) Origin of finches and diversification of the *P. pinnatus+Joubertophyllodes* clade.

The frequencies of residuals produced by Random TaPas with GD mapped on the tanglegram indicated higher congruence between the Passeroidea and their associated mite lineages than between the other clades of birds and mites (Fig. 9). Similar results were obtained applying Random TaPas with PACo (Fig. S8, Supplementary Material).

### Example 2: Neotropical orchids and their euglossine pollinators

The distributions of residuals (observed – expected frequencies) of the pollinator-orchid associations produced by Random TaPas with GD and PACo and their 95% confidence intervals derived from the comparison of 1,000 pairs of posterior probability trees are presented in Figure 10. Only in 12 and 35 of a total of 129 host-symbiont associations evaluated with GD and PACo, respectively, the 95% confidence intervals included only positive values. The *G*s* of these frequencies obtained with the consensus trees were 0.768 and 0.751 for Random TaPas run with GD and PACo, respectively. These values were within the range of *G*s* obtained with the 1,000 pairs of posterior probability trees (Fig. 10c). All values of *G\** produced with GD and PACo were larger than the ⅔ threshold proposed to indicate random contributions of congruence by the host-symbiont associations. The frequencies of residuals produced with GD mapped on the tanglegram indicated no clear pattern of congruent associations being associated with particular clades and only association of the early divergent euglossine *Exaerete* sp. with a representative of *Gongora* stands out as extremely incongruent (Fig. 11). (Very similar results were obtained with PACo as shown in Figure S9, Supplementary Material.)

**Figure 10.**
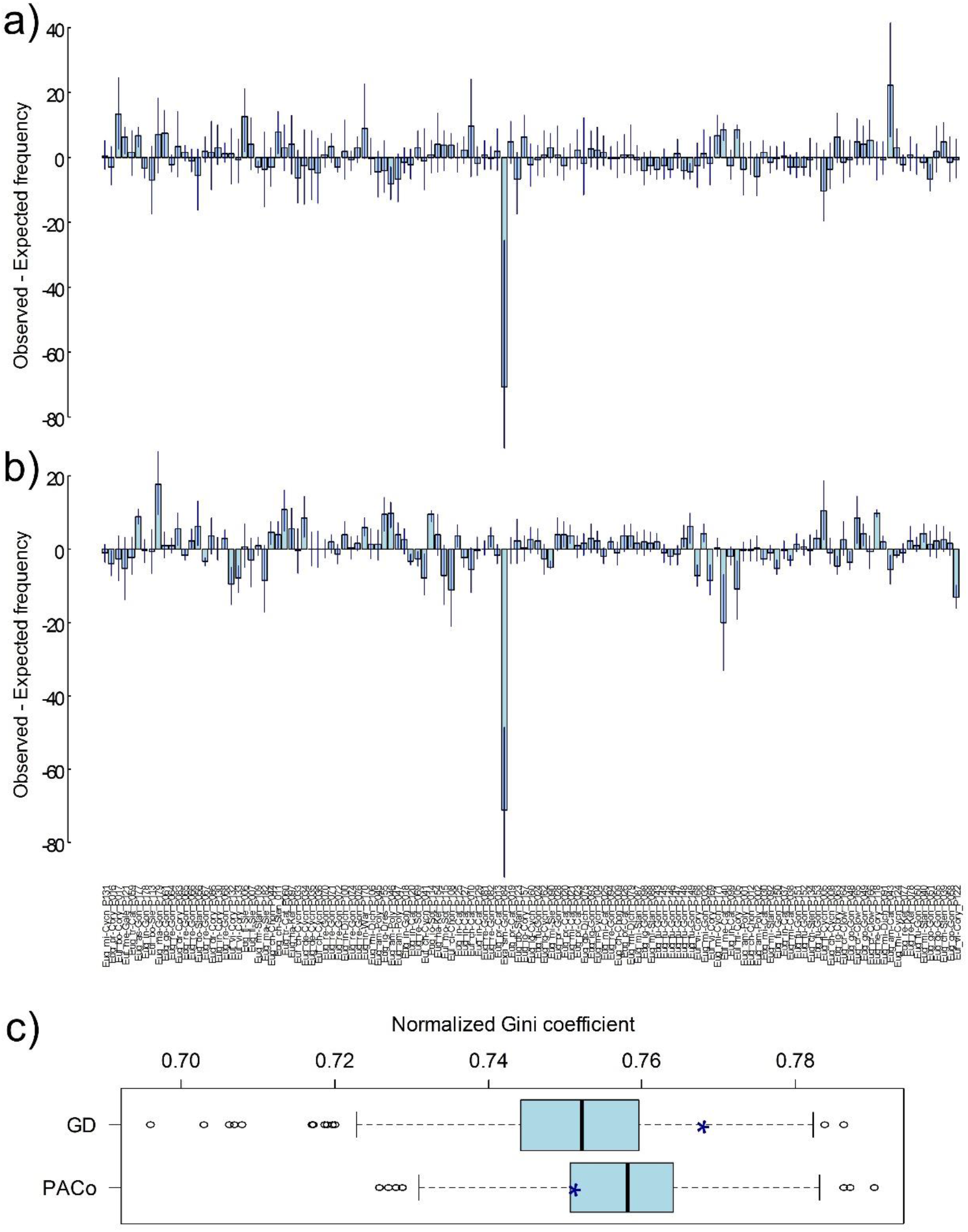
Random TaPas applied to data of Neotropical orchids and their euglossine bee pollinators. The distributions of observed – expected frequencies were obtained with *N* = 10^4^, *n* = 26, and *p* = 1% with GD (a) and PACo (b). Vertical lines represent 95% confidence intervals of the frequencies computed with 1,000 randomly chosen pairs of posterior probability trees used to build the consensus trees of bees and orchids. (c) Normalized Gini coefficient of the respective frequency distributions (asterisks) and boxplots produced with the pairs of posterior probability trees generated with GD and PACo.

**Figure 11.**
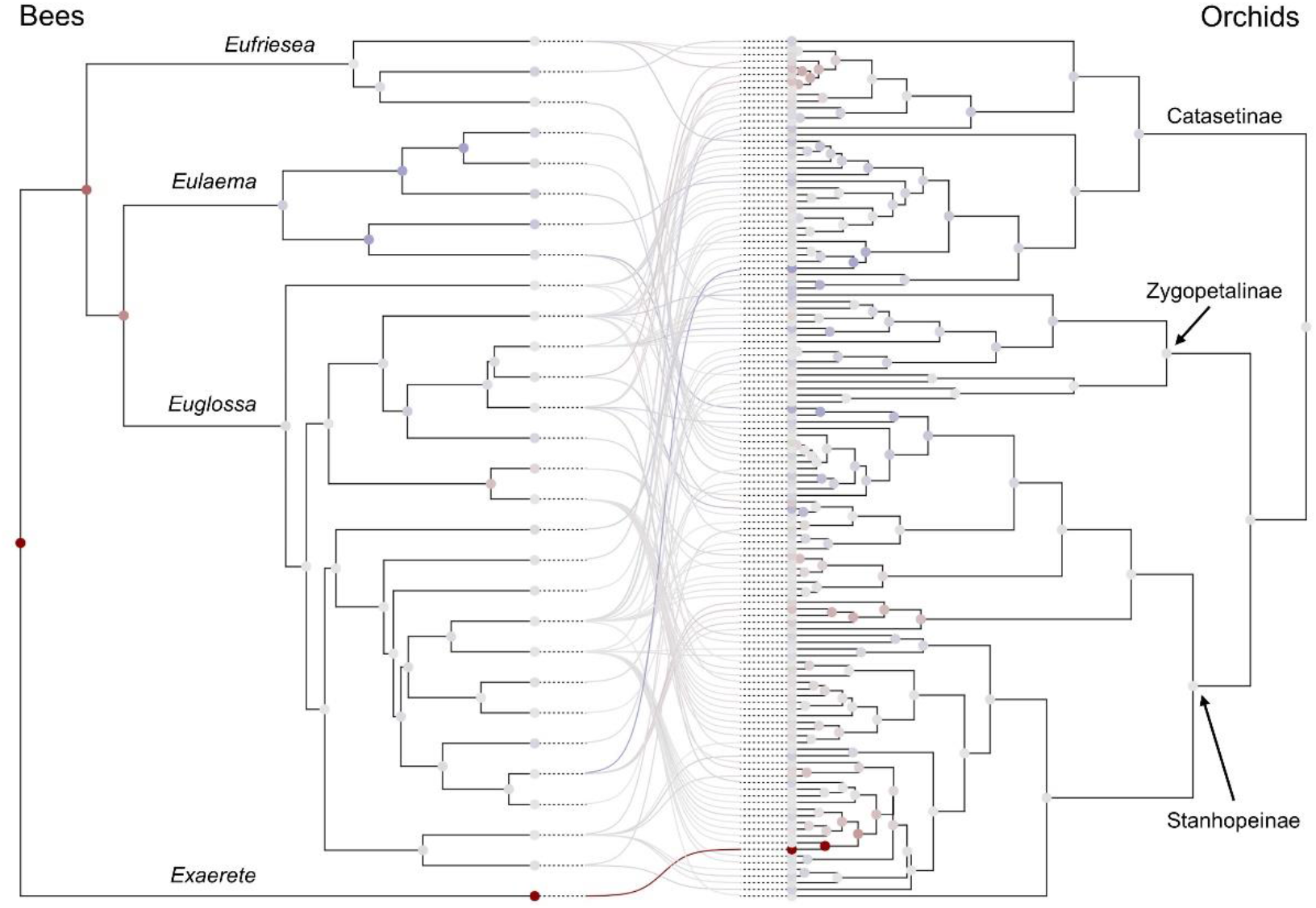
Tanglegram representing the association between Neotropical orchids and their euglossine bee pollinators. The observed-expected frequencies corresponding to each pollinator-orchid association shown in Fig. 9 obtained applying Random TaPas with geodesic distances are mapped using a diverging color scale centered at zero (light gray) and ranging from dark red to dark blue. The average observed-expected frequency of each terminal and fast maximum likelihood estimators of ancestral states of each node are also mapped according to the same scale.

## Discussion

Herein we have developed Random TaPas that uses a given global-fit method for analysis of large tanglegrams and produces a normalized Gini coefficient (*G*\*) whose ability to quantify cophylogenetic signal has been assessed by means of simulated data. Some simulations could not be run because the number of unique host-symbiont associations was less than the *n* set to perform Random TaPas (as defined in Fig. 1). This indicates that the maximum number of unique host-symbiont associations is constrained by the nature of the joint evolutionary history. Since such unique associations are potentially related to cospeciation, the maximum number that can be retrieved in a given triple can provide a first estimate of the amount of cophylogenetic signal in the system. (In the User’s Guide at GitHub, we include a function to determine the maximum *n* possible for a given triple and illustrate a strategy to set *n* optimally).

The simulation results indicated that *G*\* is a reasonable proxy for cophylogenetic signal given its negative correlation with both the number and proportion of cospeciation events, and the positive correlation with the number of coevolutionary events promoting topological incongruence. This was consistently shown with most parameter combinations and with both global-fit methods used. A noticeable exception concerned the low correlation of *G*\* with number of colonization events. Perhaps this is due to the way colonization events were simulated, by spreading the associations at the terminals of the host’s tree, rather than through the internal nodes as in the other events. In any case, the simulations suggest that the relationship between *G*\* and cophylogenetic signal is stronger in ultrametric trees and decreases with the size of the triple. Based on the simulation results, we recommend setting *n* at around 10% of the total number of associations and *p* = 1% to estimate cophylogenetic signal. However, to map congruent and incongruent associations in a given triple *n* ≈ 20% and *p* = 1 % seem to show better performance to correctly identify congruent terminals, especially those at basal positions (Figs. 6 and 7).

We have shown that simulations produced by triples involving ultrametric trees in which *G*\*s < ⅔ usually correspond to *G*\*s > ⅔ in their counterparts based in additive trees. In fact, none of the latter yielded a *G*\* < 0.599 (Table S1 in Supplementary Material). Interestingly, there was a good agreement in *G*\* values between simulations with ultrametric trees rendering *G*\*s > ⅔ and the corresponding values obtained with additive trees. All this evidence suggests that, when working with phylograms, differences in branch lengths weight more in the *G*\* value as cophylogenetic signal (mostly dictated by topological congruence) increases. This is also supported by the pseudocospeciation experiment in which the triple chosen represented a scenario of high cophylogenetic signal. The simulations with additive trees indicated that the degree of incongruence of each host-symbiont association was mostly determined by the absolute differences in branch lengths of the terminals involved (Fig. 4). This would explain why associations between basal terminals tended to be marked as incongruent more often than those between recently diverged terminals (Figs. 4, 6). (This effect is also observed with ultrametric trees (Figs. 5, 7) but it is weaker.) Furthermore, the noise added by differences in branch lengths between the terminals of hosts and their symbionts could account for the lower correlations within the same set of simulated triples between *G*\* and number of coevolutionary events observed in simulations involving additive trees compared to those based on ultrametric ones.

Although we noted differences in the ranges and mean *G*\*s between Set50 and Set100, these were slight and thus *G*\* values do not seem to be critically affected by triple size (Table S1). (At least for the sizes contemplated in the present study). So for triples based on chronograms, we can propose a general framework to gauge the amount of cophylogenetic signal in any given host-symbiont system: *G*\*s between 0 and ⅔ would represent a gradient from high cophylogenetic signal, in which all host-symbiont associations would contribute similarly to the global fit between the host and symbiont phylogenies, to cophylogenetic randomness. So we predict that in systems in which *G*\* ≈ ⅔ global fit tests would often indicate no significant relationship between the evolutionary histories of hosts and symbionts. Finally, *G*\*s between ⅔ and 1 would define a gradient from random association to low cophylogenetic signal in which few host-symbiont associations would contribute mostly to the congruence or incongruence patterns observed.

However, this framework is not applicable to triples based on phylograms, since only when *G*\*s is about larger than about 0.72 we could confidently conclude that cophylogenetic signal is low (Fig. 2). However, Random TaPas is still useful as it has proven to be efficient to map congruent/incongruent associations on tanglegrams (Fig. 6). Note also that the additive trees were generated by adding varying rates of molecular substitution in the host tree independently from the corresponding symbiont tree, whereas corresponding branches in real phylograms may show some degree of dependency in their diversification rates. This is particularly evident across the plant kingdom, in which groups exhibiting very different life histories (e.g. holoparasitism, epiphytism) are linked to having disparate rates of molecular evolution and thus random, local molecular clocks explain better their mode of diversification (Bellot and Renner 2014; Wicke et al. 2016; Dann et al. 2017). This calls for additional work to account for differences in substitution rates when comparing gene trees. However, assessing temporal congruence between the phylogenies is necessary to attain a high level of confidence about the amount of cophylogenetic signal in a given system (de Vienne et al. 2007).

The quantitative framework for dated trees proposed herein is particularly valuable because current global-fit methods test for congruence between phylogenies, but do not quantify in a meaningful way the strength of the relationship because the statistics produced are not bounded and depend on the scale (branch units) of the distances computed between the terminals. For instance, in Example 1, PACo and ParaFit point to highly significant evidence for cophylogenetic signal between each of the 200 bird trees and the mite phylogeny (*P* < 0.001 in all tests). Likewise, applying the same analyses to Example 2 leads to a similar conclusion PACo (*m*^2^_*XY*_ = 9.35 10^6^, *P* < 0.0001) and ParaFit (ParaFitGlobal = 5.14·10^8^, *P* < 0.017). However, the mapping of residual frequencies on both tanglegrams suggest very different coevolutionary histories in each system. In addition, the *G*\* values in Example 1 were smaller and slightly above the ⅔ threshold, which would lead to conclude that cophylogenetic signal is low to moderate. This agrees with the results of the event-based reconciliation analysis carried out by Klimov et al. (2017) with the same dataset that yielded 52 cospeciation events out of 194 coevolutionary events. In Example 2, the high *G*\* values point to low cophylogenetic signal. After considering phylogenetic uncertainty, the confidence intervals of few residuals did not include zero, and congruent and incongruent associations were scattered in the tanglegram, without showing a clear pattern. All this evidence suggests that that cospeciation events were not the main driver accounting for the coevolution of Neotropical orchids and their euglossine pollinator bees. This conclusion is in line with the hypothesis that preexisting traits in the euglossine bees (i.e., collection of aromatic compounds), rather than cospeciation, drove floral adaptation and diversification in the euglossine bee pollinated orchids (Ramírez et al. 2011; Pérez-Escobar et al. 2017). In contrast, in Example 1, cophylogenetic signal was unevenly distributed being stronger between the Passeroidea and their associated feather mites. This topological agreement reflects well the synchronic diversification at about 20 Mya of two passeroid sister clades, the New World emberizoids and Old World finches, with two corresponding sister clades of feather mites, the *Proctophyllodes thraupis+quadratus* and *P. pinnatus+Joubertophyllodes* clades (Klimov et al. 2007) (Fig. 10).

Random TaPas has been developed with two very different global-fit approaches. As noted above a potential of GD over PACo, and most other global-fit methods, is that it skips the potential effect of the dimensional mismatch between Euclidean and tree spaces (Holmes 2005). However, GD performed only marginally better than PACo in terms of relationship between *G*\* and number of cospeciation events, and overall results were very similar. This evidence points to robustness of Random TaPas to parameters and method utilized. In fact, the algorithm can be readily adapted to other global-fit methods. In a preliminary analysis, we tried it with ParaFit (Legendre et al. 2002) and some parameter combinations yielding very similar results to those presented here in terms of relationship with the number of cospeciation events (Table S2, Supplementary Material).

A problem with global-fit methods is that they do not correct for phylogenetic nonindependence, so that comparison of pairwise distance matrices derived from host and symbiont phylogenies gives greater weight to pairs that include deeper most recent common ancestors (MRCAs) than those involving shallower ones (Schardl et al. 2008; de Vienne et al. 2013). Without correction for nonindependence, phylogenetic congruence between old MRCAs is expected to be propagated towards the descendent terminals. Thus associations between them would tend to be identified as congruent even if the placement of their more recent MRCAs is not. This results in distance-based tests being anticonservative when evaluating the contribution of individual links to the global fit. To deal with this issue, Schardl et al. (2008) proposed selecting a single pairwise distance per MRCA prior to evaluation of congruence between two phylogenies globally. However, this approach is not immediately applicable to assess the contribution of individual host-symbiont associations to the global fit.

So although further work is needed, we suggest that, even though nonindependence is not completely accounted for in the partial tanglegrams, the evaluation of a limited set of associations over a large *N* seems to buffer the effect of phylogenetic nonindependence. So Random TaPas would represent a better way to determine which host-symbiont associations contribute most to cophylogenetic signal than current global-fit methods. In any event, we find that the mapping of cophylogenetic signal on the tanglegram is an extremely useful new tool for the analysis of coevolutionary histories as it allows evaluating variation in cophylogenetic signal and testing specific hypotheses.

Interpretation of the results produced by Random TaPas is not without some issues. A known limitation of the Gini coefficient is that different patterns of inequality can yield the same value of the coefficient (Ultsch and Löstch, 2017). So users may need to examine the Lorenz curves associated to each sample in order to discriminate systems with similar G*s. (Raffinetti et al. (2015) describe a procedure to derive a generalized Lorenz curve from data including negative attributes.) Future studies would also need to explore the performance of alternative metrics of inequality, such as the Atkinson index or the Generalized Entropy index, but this requires modifications to allow the incorporation of negative observations.

One should also bear in mind that host-symbiont associations are marked as congruent or incongruent relative to the other associations in the triple. As indicated above, this is more apparent when working with phylograms but the issue also affects chronograms. Since Random TaPas is based on distance-based algorithms, the absolute differences in branch lengths between pairs of host-symbiont associations influences the estimation of the associated residual value. So, everything else being equal, a basal host-symbiont association is more likely to have a lower residual than more distal ones. By the same token, incongruent associations in upper branches can be scored as congruent. In Example 1, for instance, the associations between *Emberiza* spp., embedded within the New World emberizoids, and the *P. pinnatus+Joubertophyllodes* clade is indicative of host-switches from finches, but despite this they were plotted as congruent on the tanglegram (Figure 12). A possible way to tackle this problem could be to run partial analyses with designated nodes, for instance, within the Passeroidea and their mites in order to get a more detailed picture at a lower taxonomic level. Hoyal-Cuthill and Charleston (2012, 2015) adopted a similar approach and we consider that its potential integration with Random TaPas is worth exploring in the future.

A final word of caution is that the effects of cospeciation and pseudocospeciation on cophylogenetic signal can be difficult to tease apart, even if the trees are dated. However, one would expect pseudocospeciation to be more prevalent if host trees are clustered in few large clades and in those with rapid species turnover (rapid adaptive radiations), particularly if symbionts are highly specific (Engelstädter and Fortuna 2019). With dated trees, our results suggest that marked differences in frequency of host-symbiont associations with varying *n* can give clues about the presence of pseudocospeciation in the system.

In summary, Random TaPas represents a new tool for cophylogenetic analysis that provides a framework to assess cophylogenetic signal in a given host-symbiont system. In addition, it facilitates data interpretation by mapping the extent of cophylogenetic signal on the tanglegram. The method can handle large tanglegrams in affordable computational time, incorporates phylogenetic uncertainty, makes a more explicit links with cospeciation and other coevolutionary events, and is more reliable to identify host-symbiont associations that contribute most to cophylogenetic signal than regular global-fit methods. For greater usability, Random TaPas is implemented in the public-domain statistical software R.

## Supporting information

Supp. Tables, Figures and proof.

## Acknowledgements

We are extremely grateful to Santiago Ramírez, University of California Davis, and to Pavel Klimov, University of Michigan, for generously providing the raw data of the real-world examples tested and for their valuable indications for the analysis and interpretation of results.

## References

Balbuena J.A., Míguez-Lozano R., Blasco-Costa I. 2013. PACo: A novel Procrustes application to cophylogenetic analysis. PLoS ONE 8(4):e61048.

Bellot S., Renner S.S. 2014. Exploring new dating approaches for parasites: the worldwide Apodanthaceae (Cucurbitales) as an example. Mol. Phylogenet. Evol. 80:1–10.

Brown R.P., Yang Z. 2011. Rate variation and estimation of divergence times using strict and relaxed clocks. BMC Evol. Biol. 11:271.

Chakerian J., Holmes S. 2010. Computational tools for evaluating phylogenetic and hierarchical clustering trees. arXiv:1006.1015.

Charleston M.A., Robertson D.L. 2002. Preferential host switching by primate lentiviruses can account for phylogenetic similarity with the primate phylogeny. Syst. Biol. 51:528–535.

Charleston M.A., Libeskind-Hadas R. 2014. Event-based cophylogenetic comparative analysis. In: Garamszegi L.Z., editor. Modern phylogenetic comparative methods and their application in evolutionary biology. Berlin, Heidelberg: Springer. p. 465–480.

Dann M., Bellot S., Schepella S., Schaefer H., Tellier A. 2017. Mutation rates in seeds and seed-banking influence substitution rates across the angiosperm phylogeny. bioRxiv: 156398.

de Vienne D.M., Aguileta G., Ollier S. 2011. Euclidean nature of phylogenetic distance matrices. Syst. Biol. 60:826–832.

de Vienne D.M., Giraud, D T., Shykoff J.A. 2007. When can host shifts produce congruent host and parasite phylogenies? A simulation approach. J. Evol. Biol. 20:1428–1438.

de Vienne D.M., Refrégier G., López-Villavicencio M., Tellier A., Hood M.E., Giraud T. 2013. Cospeciation vs host-shift speciation: Methods for testing, evidence from natural associations and relation to coevolution. New Phytol. 198:347–385.

Drinkwater B., Qiao A., Charleston M.A. 2016. WiSPA: A new approach for dealing with widespread parasitism. arXiv:1603.09415

Engelstädter J., Fortuna N.Z. 2019. The dynamics of preferential host switching: Host phylogeny as a key predictor of parasite distribution. Evolution 73-7:1330–1340.

Hagen O., Hartmann K., Steel M., Stadler T. 2015. Age-dependent speciation can explain the shape of empirical phylogenies. Syst. Biol. 64:432–440.

Holmes S. 2005. Statistical approach to tests involving phylogenies. In: Gascuel O., editor. Mathematics of Evolution and Phylogeny. Oxford: Oxford University Press. p. 91–120.

Hoyal-Cuthill J., Charleston M. 2012. Phylogenetic codivergence supports coevolution of mimetic *Heliconius* butterflies. PLoS ONE 7(5):e36464.

Hoyal-Cuthill J., Charleston M. 2015. Wing patterning genes and coevolution of Müllerian mimicry in *Heliconius* butterflies: Support from phylogeography, cophylogeny, and divergence times. Evolution 69:3082–3096.

Hutchinson M.C., Cagua E.F., Balbuena J.A., Stouffer D.B., Poisot T. 2017a. paco: implementing Procrustean Approach to Cophylogeny in R. Meth. Ecol. Evol. 8:932–940.

Hutchinson M.C., Cagua E.F., Stouffer D.B. 2017b. Cophylogenetic signal is detectable in pollination interactions across ecological scales. Ecology. 98:2640–2652.

Kahnt B., Hatting W.N., Theodorou P., Wieseke N., Kuhlmann M., Glennon K.L., van der Niet T., Paxton R., Cron G.V. 2019. Should I stay or should I go? Pollinator shifts rather than cospeciation dominate the evolutionary history of South African *Rediviva* bees and their *Diascia* host plants. Mol. Ecol. in press. DOI: 10.1111/mec.15154.

Keller-Schmidt S., Wieseke N., Klemm K., Middendorf M. 2011. Evaluation of host parasite reconciliation methods using a new approach for cophylogeny generation. Technical Report, Bioinformatics Leipzig. Available at http://www.bioinformatik.uni-leipzig.de/Publications/PREPRINTS/11-013.pdf

Klimov P.B., Mironov S.V., OConnor B.M. 2017. Detecting ancient codispersals and host shifts by double dating of host and parasite phylogenies: Application in proctophyllodid feather mites associated with passerine birds. Evolution 71: 2381–2397.

Lagrue C., Joannes A., Poulin R., Blasco-Costa I. 2016. Genetic structure and host – parasite co-divergence: evidence for trait-specific local adaptation. Biol. J. Linn. Soc. 118:344–358.

Legendre P., Desdevises Y., Bazin E. 2002. A statistical test for host-parasite coevolution. Syst. Biol. 51:217–234.

Mendlová M., Desdevises Y., Civáňová K., Pariselle A., Šimková A. 2012. Monogeneans of West African cichlid fish: evolution and cophylogenetic interactions. PLoS ONE 7(5):e37268.

Moran N.A. 2006. Symbiosis. Current Biol. 16:R866–R871.

Page R.D.M. 2003. Introduction. In: Page R.D.M., editor. Tangled trees: phylogeny, cospeciation and coevolution. Chicago: Chicago University Press. p. 1–22.

Paradis E. 2014. Simulation of phylogenetic data. In: Garamszegi L.Z., editor. Modern phylogenetic comparative methods and their application in evolutionary biology. Berlin, Heidelberg: Springer Verlag. p. 335–350.

Pérez-Escobar O.A., Balbuena J.A., Gottschling M. 2016. Rumbling orchids: How to assess divergent evolution between chloroplast endosymbionts and the nuclear host. Syst. Biol. 65:51–65.

Pérez-Escobar O.A., Chomicki G., Condamine F.L., Karremans A., Bogarin D., Matzke N.J., Silvestro D, Antonelli A. 2017. Recent origin and rapid speciation of Neotropical orchids in the world’s richest plant biodiversity hotspot. New Phytol. 215:891–905.

R Core Team. 2018. R: A language and environment for statistical computing. R Foundation for Statistical Computing, Vienna, Austria.

Raffinetti E., Aimar F. 2016. Computing the Gini-based coefficients for weighted and negative attributes. https://cran.r-project.org/web/packages/GiniWegNeg/GiniWegNeg.pdf.

Raffinetti E., Siletti E., Vermizzi A. 2015. On the Gini coefficient normalization when attributes with negative values are considered. Stat. Methods Appl. 24:507–521.

Ramírez S.R., Eltz T., Fujiwara M.K., Gerlach G., Goldman-Huertas B., Tsutsui N.D., Pierce N.E. 2011. Asynchronous diversification in a specialized plant-pollinator mutualism. Science 333:1742–1746.

Revell L.J. 2012. phytools: An R package for phylogenetic comparative biology (and other things). Meth. Ecol. Evol. 3:217–223.

Schardl C.L., Craven K.D., Speakman S., Stromberg A., Lindstrom A., Yoshida R. 2008. A novel test for host-symbiont codivergence indicates ancient origin of fungal endophytes in grasses. Syst. Biol. 57:483–498.

Ultsch A., Lötsch J. 2017. A data science based standardized Gini index as a Lorenz dominance preserving measure of the inequality of distributions. PLoS ONE 12(8):e0181572.

Venables W.N., Ripley B.D. 2002. Modern applied statistics with S. New York: Springer.

Weber M.G., Wagner C.E., Best R.J., Harmon L.J., Matthews B. 2017. Evolution in a community context: On integrating ecological interactions and macroevolution. Trends Ecol. Evol. 32:291–304.

Wicke S., Müller K.F., dePamphilis C.W., Quandt D., Bellot S., Schneeweiss S.M. 2016. Mechanistic model of evolutionary rate variation en route to a nonphotosynthetic lifestyle in plants. PNAS 113:9045–9050.

Zook D. 2015. Symbiosis – evolution’s co-author. In: Gontier N., editor. Reticulate evolution. Cham, Switzerland: Springer. p. 41–80.

