## Supplementary material for "Random Tanglegram Partitions (Random TaPas): An Alexandrian Approach to the Cophylogenetic Gordian Knot": Supp. Tables, Figures and proof.

### NORMALIZED GINI COEFFICIENT OF A UNIFORM RANDOM DISTRIBUTION CENTERED AT ZERO

Here we prove that the normalized Gini coefficient ($G^*$) proposed by Raffinetti et al. (2015) of a continuous random distribution centered at zero is ⅔.

The Gini coefficient ($G$) can be calculated from the cumulative distribution function ($F(x)$) as

$$\mathrm{G} = \frac{1}{\mu} \int_{-\infty}^{+\infty} F(x)\big(1 - F(x)\big)\, dx$$

where $\mu$ represents the mean of the distribution. Given that $G^*$ is computed in an analogous fashion by replacing $\mu$ with the normalized term $\mu_Y^*$, it follows that

$$\mathrm{G}^* = \frac{1}{\mu_Y^*} \int_{-\infty}^{+\infty} F(x)\big(1 - F(x)\big)\, dx \qquad (1)$$

The cumulative distribution function of a random uniform distribution bounded between $a$ and $b$ is $F(x) = \frac{x-a}{b-a}$. Since our distribution is centered at zero (i.e., $a = -b$), $F(x) = \frac{x+b}{2b}$, and substituting in (1), $G^*$ then is expressed as:

$$\mathrm{G}^* = \frac{1}{\mu_Y^*} \int_{-b}^{+b} \frac{x+b}{2b}\left(1 - \frac{x+b}{2b}\right)\, dx \qquad (2)$$

The normalized term is computed as $\mu_Y^* = (T_Y^+ + T_Y^-)/N$, where $T_Y^+ = \sum_{i=1}^{N} max(0, y_i)$ and $T_Y^- = |\sum_{i=1}^{N} min(0, y_i)|$ (i.e., the sum of positive residuals and absolute sum of negative residuals, respectively) (Raffinetti et al. 2015). If the distribution is centered at zero and symmetrical, $T_Y^+ = T_Y^-$, it follows that $\mu_Y^* = 2T_Y^+/2n$, where $2n = N$. Since $T_Y^+/n$ is the average of a uniform random distribution between 0 and $b$, $\mu_Y^* = b/2$. Integrating and substituting in (2)

$$\mathrm{G}^* = \frac{1}{b/2} \cdot \frac{b}{3} = \frac{2}{3}$$

### REFERENCE

Raffinetti E., Siletti E., Vermizzi A. 2015. On the Gini coefficient normalization when attributes with negative values are considered. Stat. Methods Appl. 24:507-521.

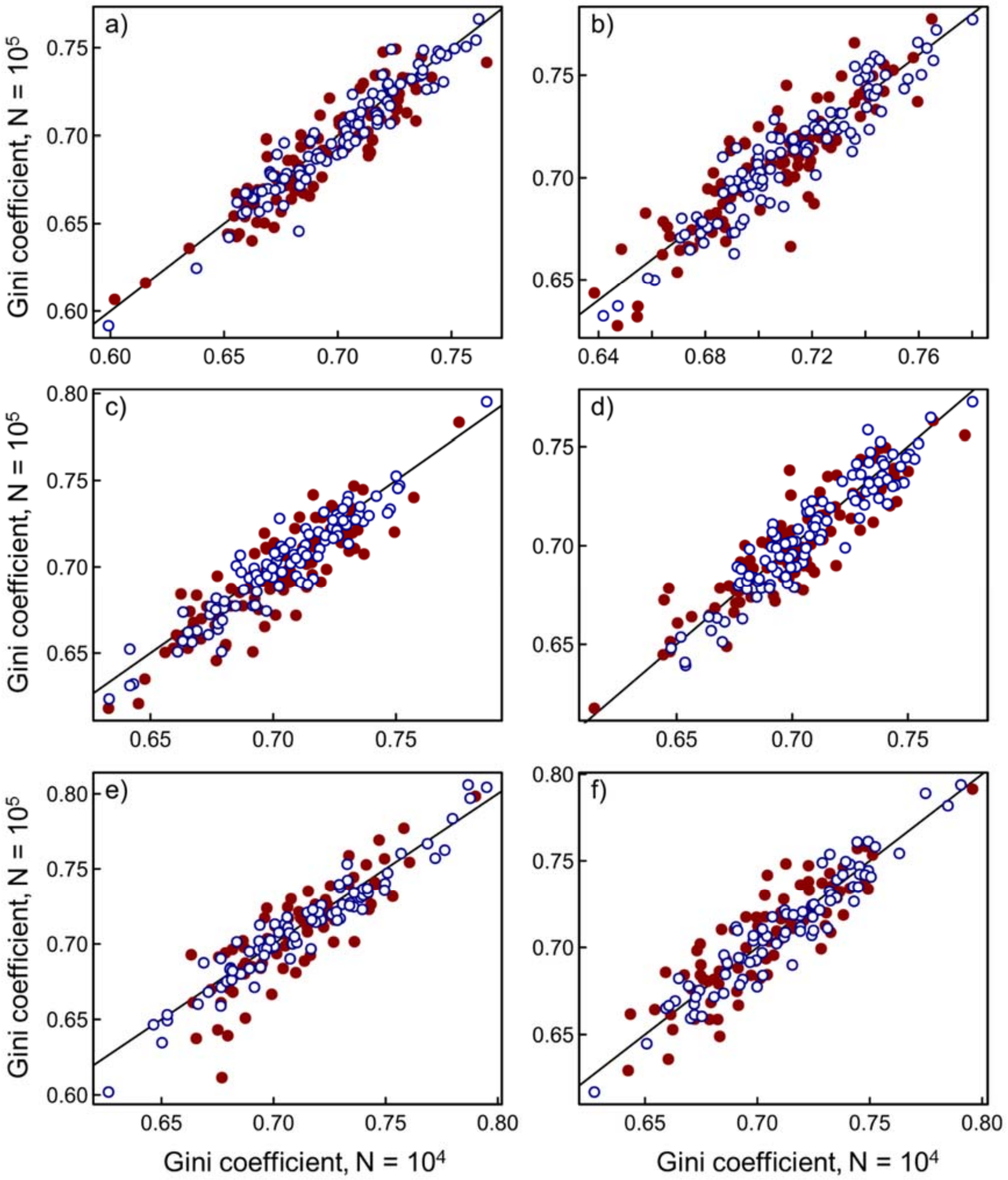


**Figure S1.** Comparison of the normalized Gini coefficients of the residual frequency distributions of host-symbiont associations produced by Random TaPas applied to 100 triples of Set50, additive trees, with numbers of runs, $N = 10^4$ vs. $N = 10^5$. Parameter and test combinations: (a) Geodesic distances (GD), $n = 5$; (b) PACo, $n = 5$; (c) GD, $n = 10$; (d) PACo, $n = 10$; (e) GD, $n = 20$; (f) PACo, $n = 20$. Filled red points, $p = 1\%$; blue empty points, $p = 5\%$. The line represents $y = x$

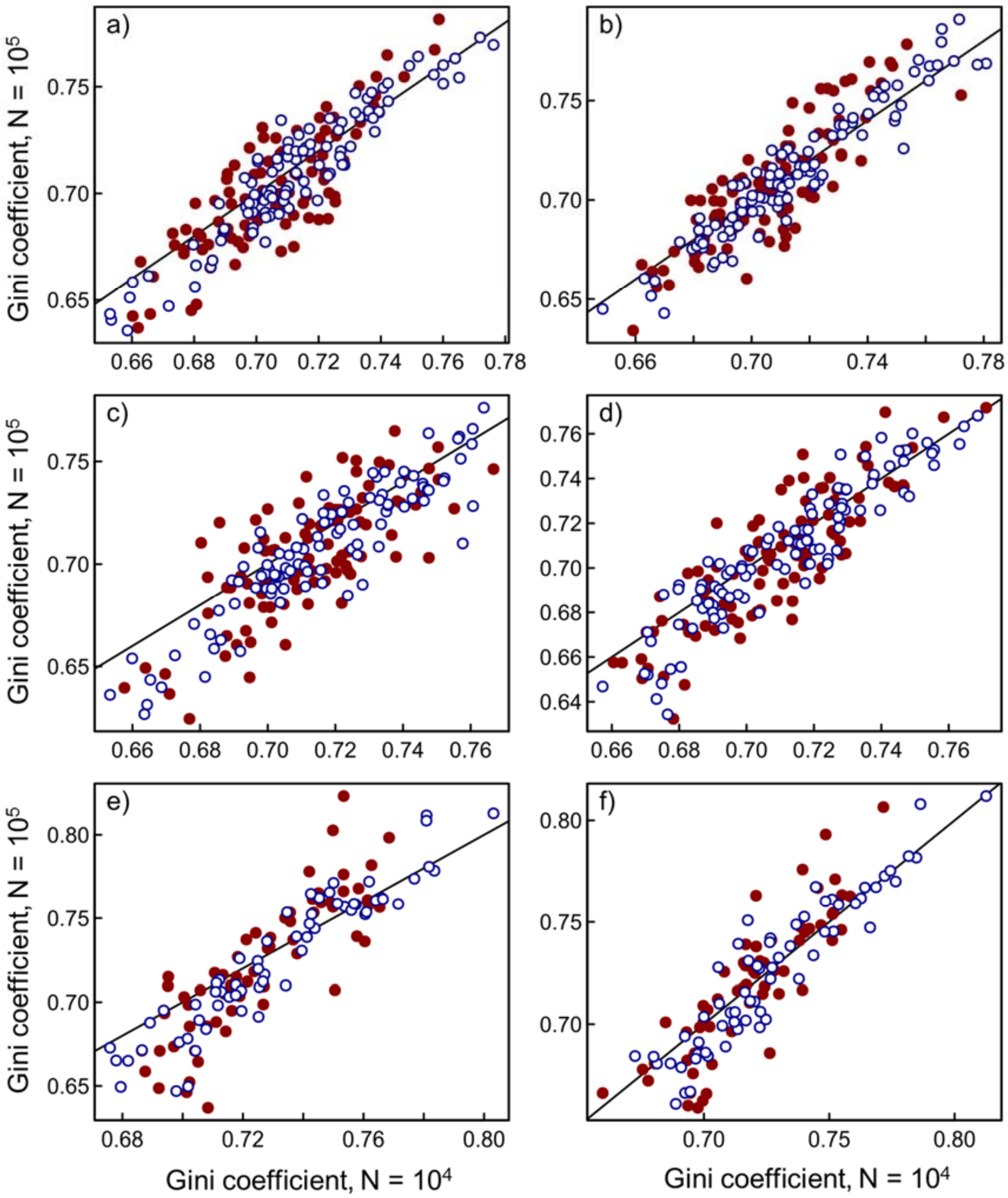


**Figure S1.** Comparison of the normalized Gini coefficients of the residual frequency distributions of host-symbiont associations produced by Random TaPas applied to 100 triples of Set100, additive trees, with numbers of runs, $N = 10^4$ vs. $N = 10^5$. Parameter and test combinations: (a) Geodesic distances (GD), $n = 10$; (b) PACo, $n = 10$; (c) GD, $n = 20$; (d) PACo, $n = 20$; (e) GD, $n = 40$; (f) PACo, $n = 40$. Filled red points, $p = 1\%$; blue empty points, $p = 5\%$. The line represents $y = x$

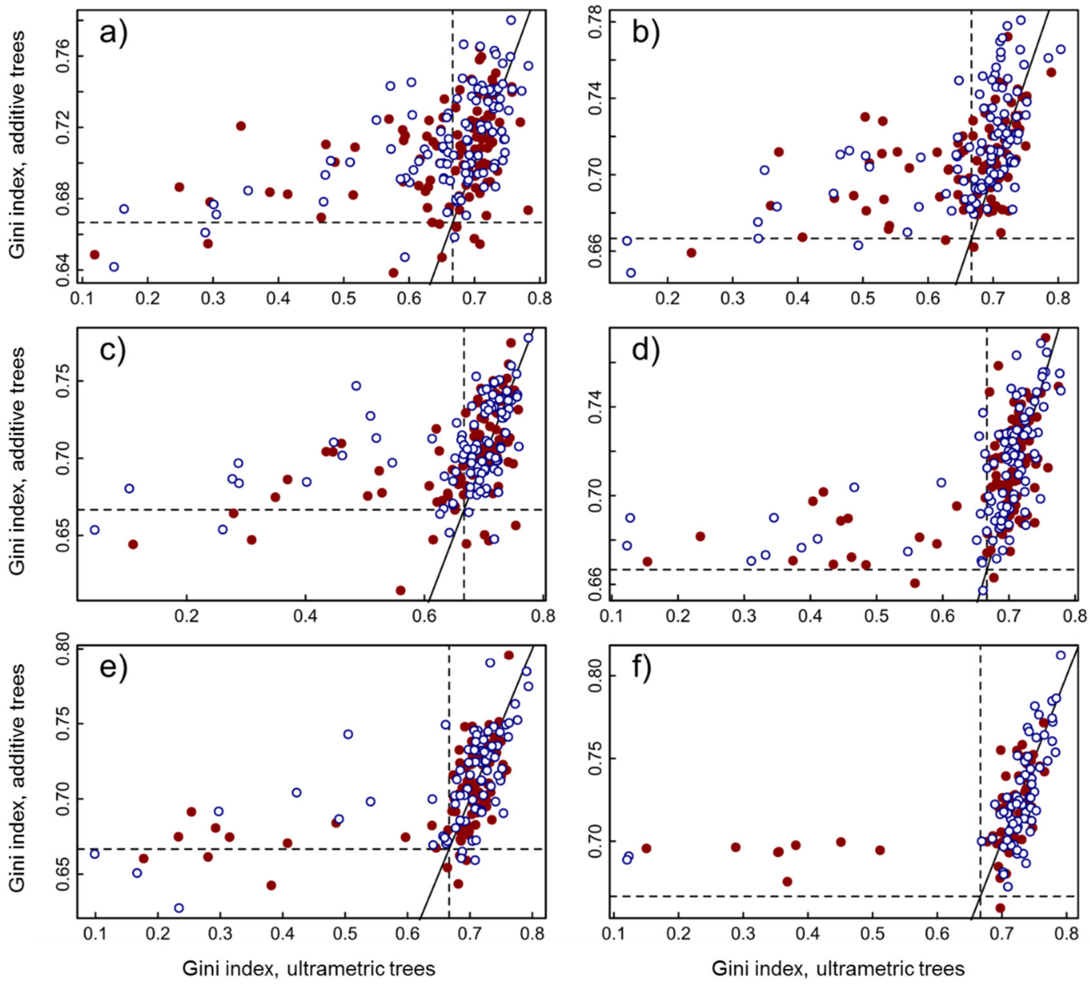


**Figure 3S.** Comparison of the normalized Gini coefficient of the residual (observed - expected frequency) distribution of host-symbiont associations produced by Random TaPas using PACo as global-fit method and ultrametric trees, with those based on additive trees. Parameter and test combinations: (a) Set50, $n = 5$; (b) Set100, $n = 10$; (c) Set50, $n = 10$; (d) Set100, $n = 20$; (e) Set50, $n = 20$; (f) Set100, $n = 40$. Filled red points, $p = 1\%$; empty blue points, $p = 5\%$. The dashed lines mark a theoretical threshold (⅔) between low and high cophylogenetic signal. The solid line represents $y = x$.

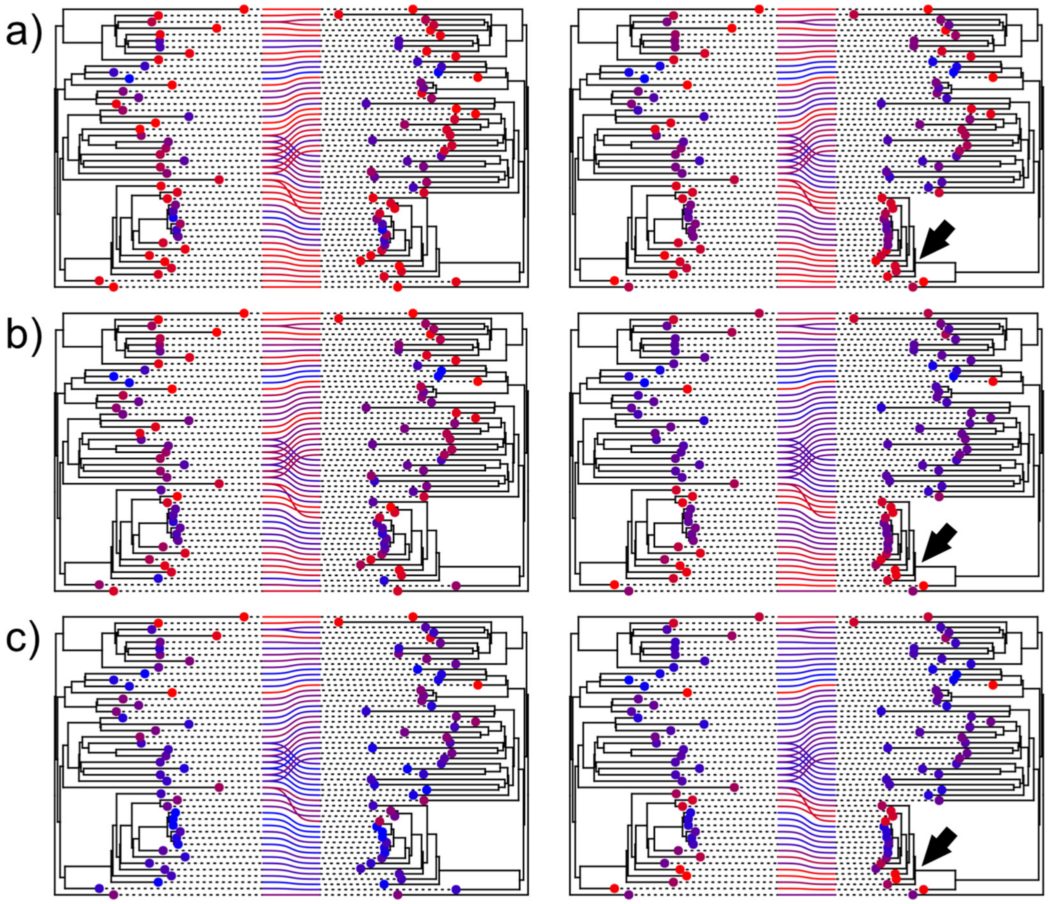


**Figure S4.** Pseudocospeciation experiment with one simulated tanglegram of ≈50 host-symbiont associations relating two additive trees. Random TaPas was applied with PACo, $p$ = 1% and $n$ = 5 (a), $n$ = 10 (b) and $n$ = 20 (c) to the original (left) and modified (right) tanglegrams. In the latter, the branch lengths of one clade (arrow) were reduced to one half, whereas its basal branch was lengthened to keep the original height of the clade. The residual (difference between observed and expected frequency of occurrence of each host-symbiont association) in the percentile $p$ retrieved by Random TaPas (see Fig. 1) is coded in a color scale, where red and blue denote low and high values, respectively. The points at terminals are also coded on the same scale and represent the average residual in which the terminal is involved.

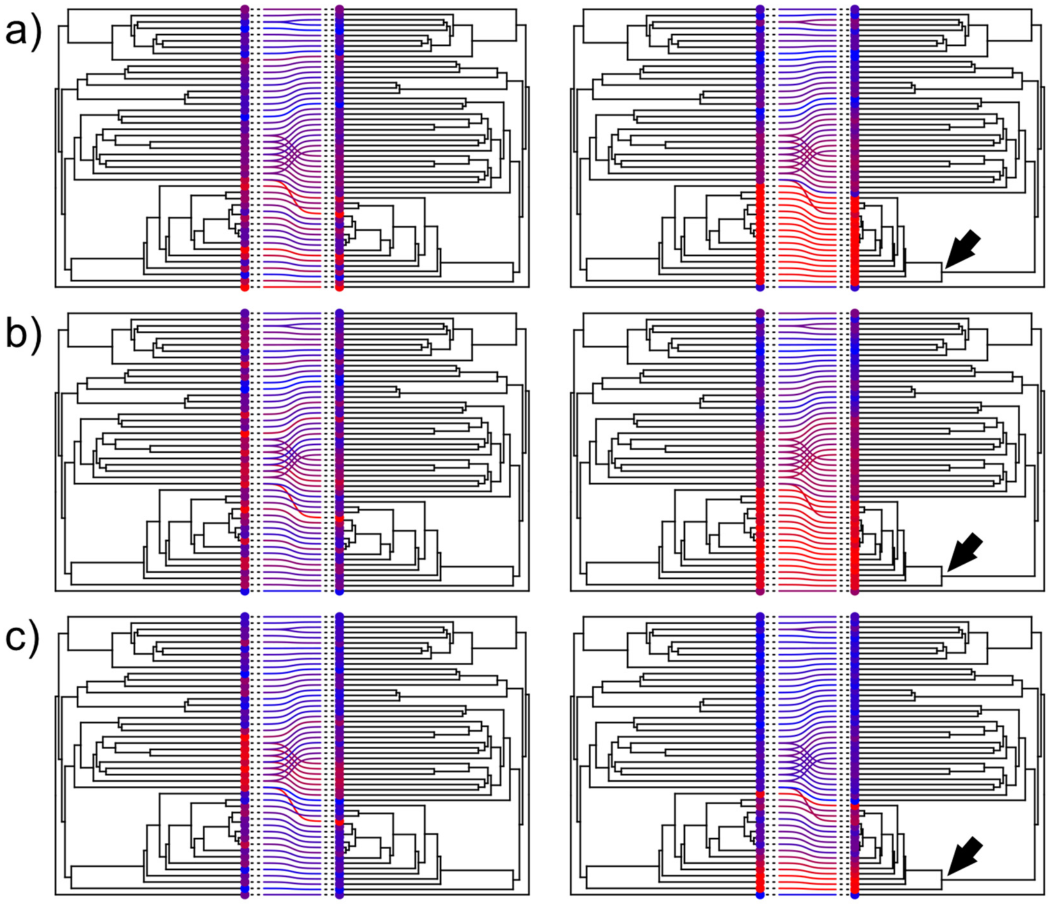


**Figure S5.** Pseudocospeciation experiment with one simulated tanglegram of ≈50 host-symbiont associations relating two ultrametric trees. Random TaPas was applied with PACo, $p$ = 1% and $n$ = 5 (a), $n$ = 10 (b) and $n$ = 20 (c) to the original (left) and modified (right) tanglegrams. In the latter, the branch lengths of one clade (arrow) were reduced to one half, whereas its basal branch was lengthened to keep the original height of the clade. The residual (difference between observed and expected frequency of occurrence of each host-symbiont association) in the percentile $p$ retrieved by Random TaPas (see Fig. 1) is coded in a color scale, where red and blue denote low and high values, respectively. The points at terminals are also coded on the same scale and represent the average residual in which the terminal is involved.

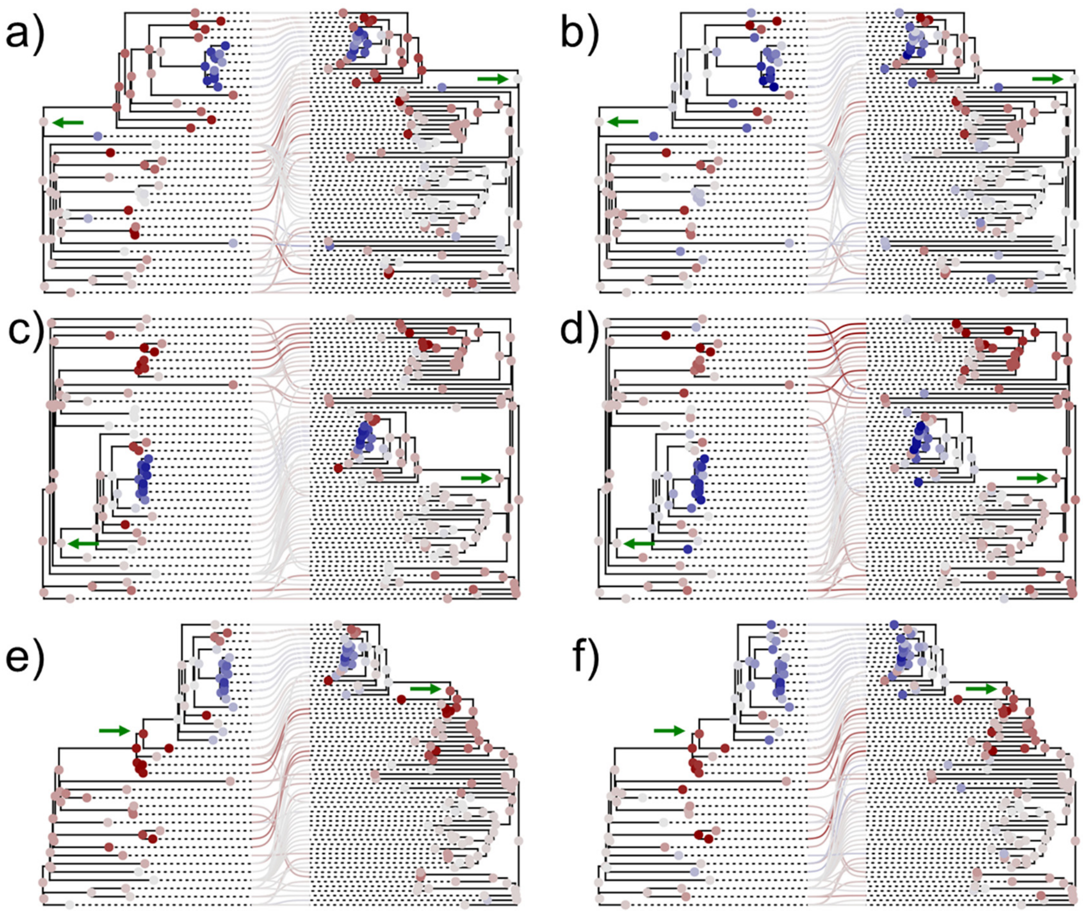


**Figure S6.** Simulated tanglegrams with additive trees in which a clade of host and symbionts (arrows) were inserted at different levels of the receptor tanglegram: at the root (a, b), and at middle (c, d) and upper (e, f) nodes. Random TaPas was applied with PACo, $p$ = 1% and $n$ = 7 (a, c, e), $n$ = 14 (b, d, f) to each tanglegram. The residual (difference between the observed and expected frequency of occurrence of each host-symbiont association) in the percentile $p$ retrieved by Random TaPas (see Fig. 1) is mapped using a diverging color scale centered at zero (light gray) and ranging from dark red (maximum negative) to dark blue (maximum positive). The average observed-expected frequency of each terminal and fast maximum likelihood estimators of ancestral states of each node are also mapped according to the same scale.

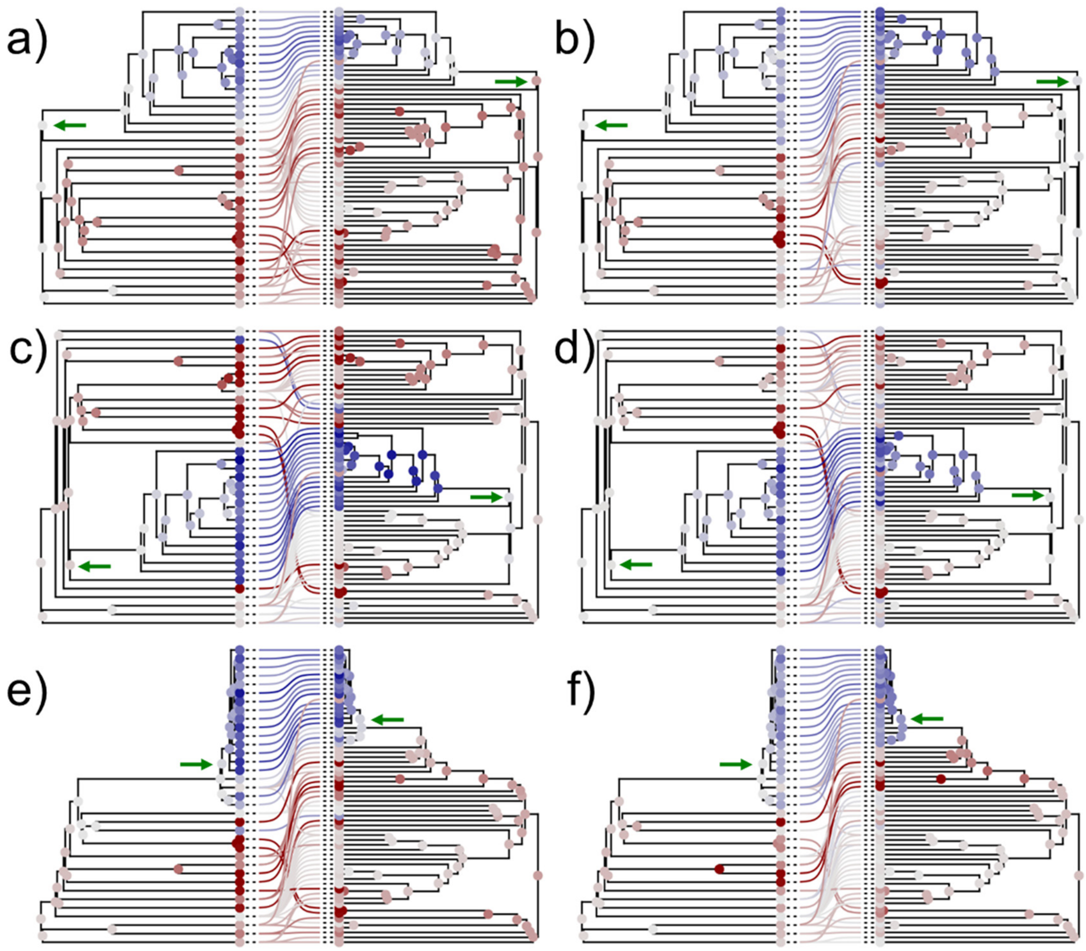


**Figure S7.** Simulated tanglegrams with ultrametric trees in which a clade of host and symbionts (arrows) were inserted at different levels of the receptor: at the root (a, b), and at middle (c, d) and upper (e,f) nodes. Random TaPas was applied with PACo, $p$ = 1% and $n$ = 7 (a, c, e), $n$ = 14 (b, d, f) to each tanglegram. The residual (difference between the observed and expected frequency of occurrence of each host-symbiont association) in the percentile $p$ retrieved by Random TaPas (see Fig. 1) is mapped using a diverging color scale centered at zero (light gray) and ranging from dark red (maximum negative) to dark blue (maximum positive). The average observed-expected frequency of each terminal and fast maximum likelihood estimators of ancestral states of each node are also mapped according to the same scale.

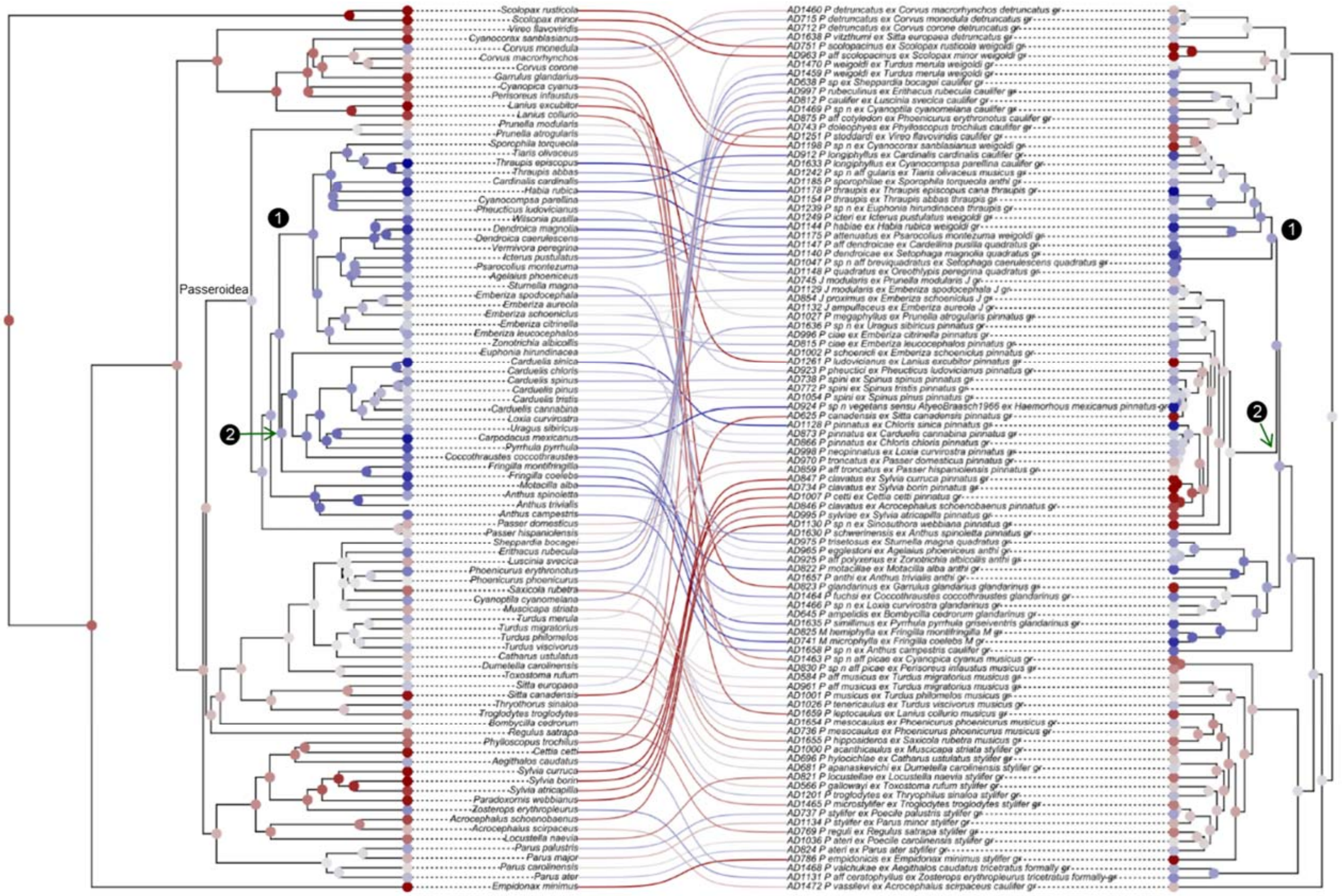


**Figure S8.** Tanglegram representing the association between passerine birds and their associated proctophyllodid feather mites. Random TaPas with PACo was applied to 200 bird chronograms and one mite dated tree. For each host tree, a separate analysis was performed yielding a vector of residuals (observed-expected frequencies). The average residual over the 200 runs corresponding to each bird-mite association is mapped using a diverging color scale centered at zero (light gray) and ranging from dark red (maximum incongruence) to dark blue (maximum congruence). The average residual at each terminal and fast maximum likelihood estimators of ancestral states of each node are also mapped according to the same scale. Based on Klimov et al. (2017), two synchronic events are also indicated on the host and mite chronograms: 1) Diversifications of New World emberizoid Passerida and the *Proctophyllodes thraupis*+*quadratus* clade and 2) Origin of finches and diversification of the *P. pinnatus*+*Joubertophyllodes* clade.

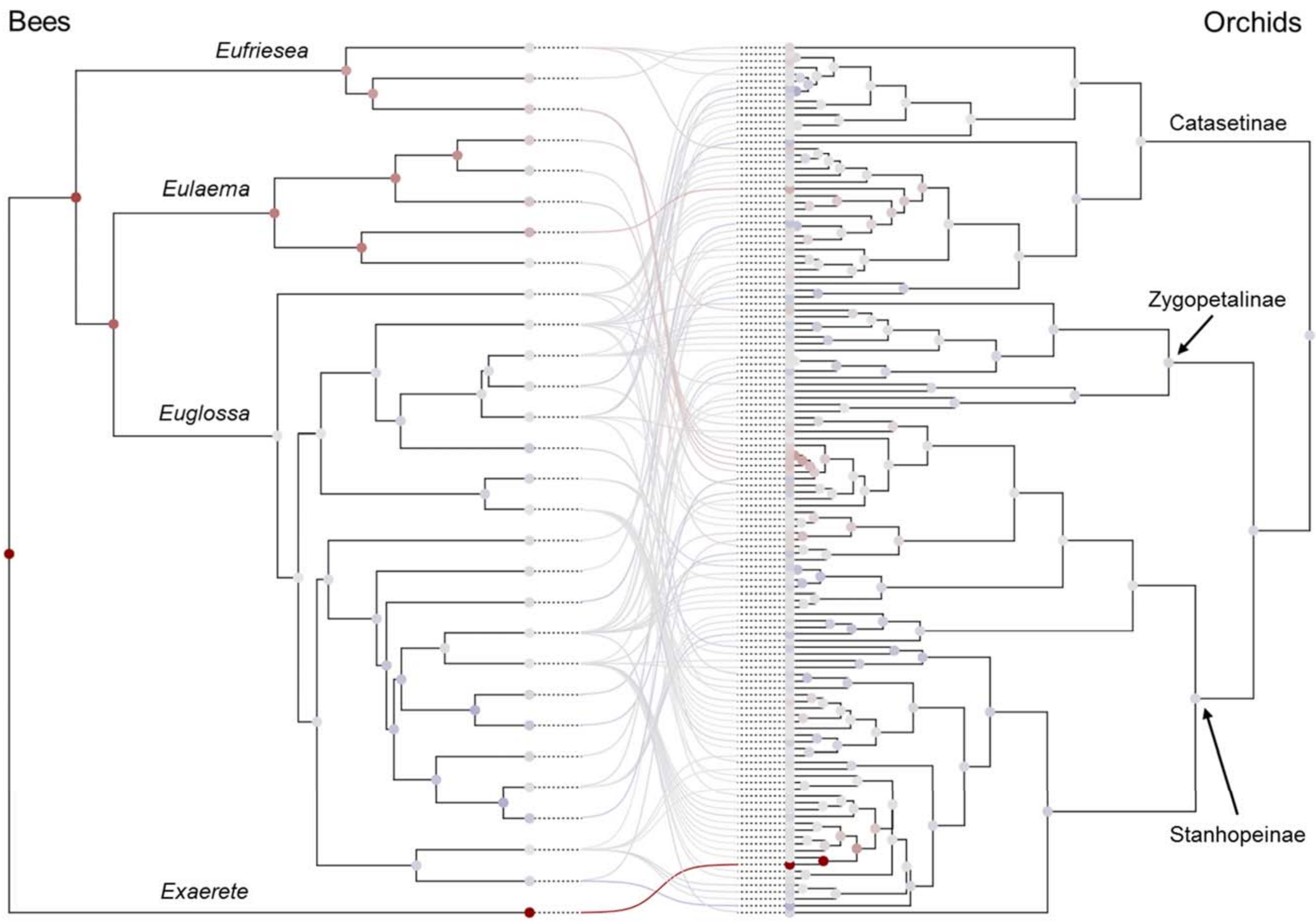


**Figure S9.** Tanglegram representing the association between Neotropical orchids and their euglossine bee pollinators. The observed-expected frequencies corresponding to each pollinator-orchid association shown in Fig. 9 obtained applying Random TaPas with PACo are mapped using a diverging color scale centered at zero (light gray) and ranging from dark red to dark blue. The average observed-expected frequency of each terminal and fast maximum likelihood estimators of ancestral states of each node are also mapped according to the same scale.

**Table S1**. Mean, standard deviation (SD) and range of normalized Gini coefficients of frequency residual distributions of host-symbiont (h-s) associations produced by Random TaPas applied in conjunction to two global-fit (GF) methods, Geodesic Distances (GD) and Procrustes Approach to Cophylogeny (PACo) to two sets of 100 simulated tanglegrams each: Set50 and Set100 involving ≈ 50 and 100 h-s associations, respectively. Random TaPas was evaluated with both additive and ultrametric trees over $10_4$ runs with a fixed number of unique h-s associations (*n*) ≈ 10%, 20% and 40% of the total number of h-s associations and percentiles (*p*) = 1% and 5%. (See Fig. 1 for definitions of *n* and *p*).

| | Set50 - Additive trees | | | Set50 - Ultrametric trees | | | Set100 - Additive trees | | | Set100 - Ultrametric trees | | |
|---|---|---|---|---|---|---|---|---|---|---|---|---|
| GF method | Mean | SD | Range | Mean | SD | Range | Mean | SD | Range | Mean | SD | Range |
| *n* ≈ 10% | | | | | | | | | | | | |
| GD, *p*=1% | 0.693 | 0.029 | 0.602 – 0.766 | 0.605 | 0.157 | 0.034 – 0.761 | 0.705 | 0.021 | 0.660 – 0.759 | 0.616 | 0.149 | 0.092 – 0.789 |
| GD, *p*=5% | 0.701 | 0.030 | 0.599 – 0.762 | 0.633 | 0.156 | 0.034 – 0.788 | 0.713 | 0.025 | 0.653 – 0.776 | 0.636 | 0.144 | 0.092 – 0.802 |
| PACo, *p*=1% | 0.704 | 0.027 | 0.638 – 0.765 | 0.649 | 0.116 | 0.119 – 0.782 | 0.706 | 0.022 | 0.659 – 0.772 | 0.660 | 0.093 | 0.237 – 0.780 |
| PACo, *p*=5% | 0.712 | 0.028 | 0.642 – 0.780 | 0.654 | 0.119 | 0.148 – 0.783 | 0.714 | 0.028 | 0.649 – 0.781 | 0.660 | 0.118 | 0.138 – 0.804 |
| *n* ≈ 20% | | | | | | | | | | | | |
| GD, *p*=1% | 0.701 | 0.026 | 0.633 – 0.776 | 0.634 | 0.150 | 0.028 – 0.764 | 0.712 | 0.021 | 0.671 – 0.757 | 0.652 | 0.143 | 0.090 – 0.767 |
| GD, *p*=5% | 0.704 | 0.027 | 0.633 – 0.787 | 0.634 | 0.151 | 0.028 – 0.789 | 0.715 | 0.026 | 0.651 – 0.767 | 0.661 | 0.136 | 0.091 – 0.763 |
| PACo, *p*=1% | 0.702 | 0.029 | 0.614 – 0.774 | 0.663 | 0.110 | 0.110 – 0.758 | 0.707 | 0.022 | 0.653 – 0.755 | 0.670 | 0.106 | 0.154 – 0.775 |
| PACo, *p*=5% | 0.708 | 0.027 | 0.648 – 0.778 | 0.655 | 0.133 | 0.045 – 0.775 | 0.712 | 0.026 | 0.653 – 0.767 | 0.671 | 0.117 | 0.123 – 0.778 |
| *n* ≈ 40% | | | | | | | | | | | | |
| GD, *p*=1% | 0.709 | 0.026 | 0.633 – 0.790 | 0.641 | 0.167 | 0.025 – 0.779 | 0.726 | 0.023 | 0.687 – 0.769 | 0.664 | 0.167 | 0.090 – 0.796 |
| GD, *p*=5% | 0.713 | 0.035 | 0.626 – 0.795 | 0.666 | 0.158 | 0.025 – 0.889 | 0.732 | 0.030 | 0.676 – 0.803 | 0.719 | 0.189 | 0.091 – 0.825 |
| PACo, *p*=1% | 0.703 | 0.028 | 0.643 – 0.796 | 0.662 | 0.132 | 0.177 – 0.763 | 0.718 | 0.024 | 0.660 – 0.712 | 0.676 | 0.129 | 0.150 – 0.766 |
| PACo, *p*=5% | 0.712 | 0.032 | 0.628 – 0.791 | 0.677 | 0.127 | 0.099 – 0.794 | 0.727 | 0.031 | 0.673 – 0.813 | 0.714 | 0.111 | 0.120 – 0.791 |

**Table S2**. Preliminary assessment of Random TaPas in conjunction with ParaFit. Correlation coefficients between normalized Gini coefficients of the frequency distribution of residuals (observed - expected frequencies) of host-symbiont associations, and the number of coevolutionary events and proportion of the number of cospeciation events with respect to the total number of events (csp/total) produced with two sets of 100 simulated triples each involving ≈ 50 and 100 host-symbiont associations, respectively, and both additive (Add) and ultrametric (Ult) trees with a varying number of host-symbiont associations (*n*) and two percentile values (*p*). (See Fig. 1 for a definition of *n* and *p*). Coevolutionary events: csp, cospeciation; sor, sorting; dup, duplication; hsw, host switching, col, colonization.

| | | | | Coevolutionary events | | | | | |
|---|---|---|---|---|---|---|---|---|---|
| Tree | Set | *p*(%) | *n* | csp | sor | dup | hsw | col | csp/total |
| Add | 100 | 1 | 10 | -0.356 | 0.327 | 0.375 | 0.249 | 0.123 | -0.399 |
| Add | 100 | 5 | 10 | -0.320 | 0.314 | 0.354 | 0.233 | 0.057 | -0.357 |
| Ult | 50 | 1 | 5 | -0.443 | 0.213 | 0.512 | 0.214 | 0.103 | -0.432 |
| Ult | 50 | 5 | 5 | -0.537 | 0.205 | 0.468 | 0.361 | 0.182 | -0.534 |
